# DeepLeMiN: Deep-learning-empowered Physics-aware Lensless Miniscope

**DOI:** 10.1101/2024.05.03.592471

**Authors:** Feng Tian, Ben Mattison, Weijian Yang

## Abstract

Mask-based lensless fluorescence microscopy is a compact, portable imaging technique promising for biomedical research. It forms images through a thin optical mask near the camera without bulky optics, enabling snapshot three-dimensional imaging and a scalable field of view (FOV) without increasing device thickness. Lensless microscopy relies on computational algorithms to solve the inverse problem of object reconstruction. However, there has been a lack of efficient reconstruction algorithms for large-scale data. Furthermore, the entire FOV is typically reconstructed as a whole, which demands substantial computational resources and limits the scalability of the FOV. Here, we developed DeepLeMiN, a lensless microscope with a custom designed optical mask and a multi-stage physics-informed deep learning model. This not only enables the reconstruction of localized FOVs, but also significantly reduces the computational resource demands and facilitates real-time reconstruction. Our deep learning algorithm can reconstruct object volumes over 4×6×0.6 mm^3^, achieving lateral and axial resolution of ∼10 µm and ∼50 µm respectively. We demonstrated significant improvement in both reconstruction quality and speed compared to traditional methods, across various fluorescent samples with dense structures. Notably, we achieved high-quality reconstruction of 3D motion of hydra and the neuronal activity with cellular resolution in awake mouse cortex. DeepLeMiN holds great promise for scalable, large FOV, real-time, 3D imaging applications with compact device footprint.

## Introduction

Fluorescence microscopy is a powerful tool in biological and biomedical research. While most of the fluorescence microscopes are in benchtop configurations, the emergence of their miniaturized counterparts enables applications demanding a compact footprint and portability, such as in endoscopic procedures or for implantable devices in bio-imaging. These miniaturized microscopes typically have the same core structure as the benchtop configurations, which incorporates an objective lens and a tube lens^1-6^. A fundamental limit of such a configuration is the tradeoff among the device footprint, field of view (FOV) and imaging resolution. Increasing the FOV while retaining high resolution tends to increase the device footprint in three dimension (3D). Furthermore, they typically lack optical sectioning and 3D imaging capability. Imaging the sample in 3D requires taking one image per focal depth by refocusing the objective lens. Mask-based lensless microscopes^7-21^, an emerging imaging technique, could overcome these limitations by replacing the bulk optics into a thin optical mask. The optical mask is in close proximity with the camera so the entire device could be very thin. The lensless microscope can resolve 3D objects in a snapshot as the optical mask can modulate the light to encode 3D object information onto a 2D image. Crucially, the FOV is readily scalable by increasing the size of the optical mask, without necessitating any increase in the device thickness. Though lensless microscope offers such advantages, it has a general challenge to recover high-quality object information. Firstly, recovering 3D information from 2D image could be an ill-posed problem, which may require strong prior information. Secondly, unlike conventional microscopes where the point spread function (PSF, describing how the object point is mapped to the camera plane) is spatially confined in a single cluster, the PSF in lensless microscopes is spatially spread out on the camera. When imaging biological samples with dense features, the images of different object points could overlap between each other. This could effectively elevate the background and thus reduce the signal contrast. Given these two reasons, even with substantial computational resources, it is challenging to reconstruct 3D volumes of dense fluorescence samples in high quality.

Here, we develop DeepLeMiN, a lensless microscope innovatively paired with a custom designed optical mask and a highly efficient physics-informed computational algorithm termed as multiFOV-ADMM-Net, enabling a high-efficient and high-quality reconstruction of both 2D and 3D objects (Fig. 1). Our optical mask features a microlens array where each unit is a lens doublet, resulting in a sparse PSF where each lobe is spatially confined. Such a PSF plays an important role to reduce the computational demand of the reconstruction algorithm. Our reconstruction algorithm distinguishes itself from others with three important features. Firstly, it is based on a multi-stage physics-informed model embedded in unrolled Alternating Direction Method of Multipliers (ADMM) deep neural networks^22,23^. Such a hybrid approach combines the advantage of the high accuracy of model-based iterative optimization approach^7,8,13,24^ and the fast running time and data adaptability of the data-driven deep neural networks^17,25,26^. Secondly, thanks to the sparse PSF which is comprised of discrete and localized lobes, our deep neural network demands considerably fewer computational resources. It could thus process data with large scale in a high speed to achieve a high reconstruction quality, which are otherwise intractable in either iterative optimization methods or deep neural networks. Thirdly, our neural network is specifically designed for imaging system that lacks a global PSF. It reconstructs the objects in local FOVs and then seamlessly integrates them together into a cohesive reconstruction over the entire FOV. DeepLeMiN has an FOV of 4×6 mm^2^, over 600 µm axial range, with an axial resolution ∼50 µm. We demonstrated high quality imaging of a variety of challenging fluorescence samples (Fig. 1), including scattering phantoms with complex and dense features, organisms like C. elegans and hydras that display 3D motion, as well as the cortical neural activity with cellular resolution in awake mouse. DeepLeMiN is a promising tool for portable, extensive field-of-view, real-time, 3D imaging applications.

**Figure 1.**
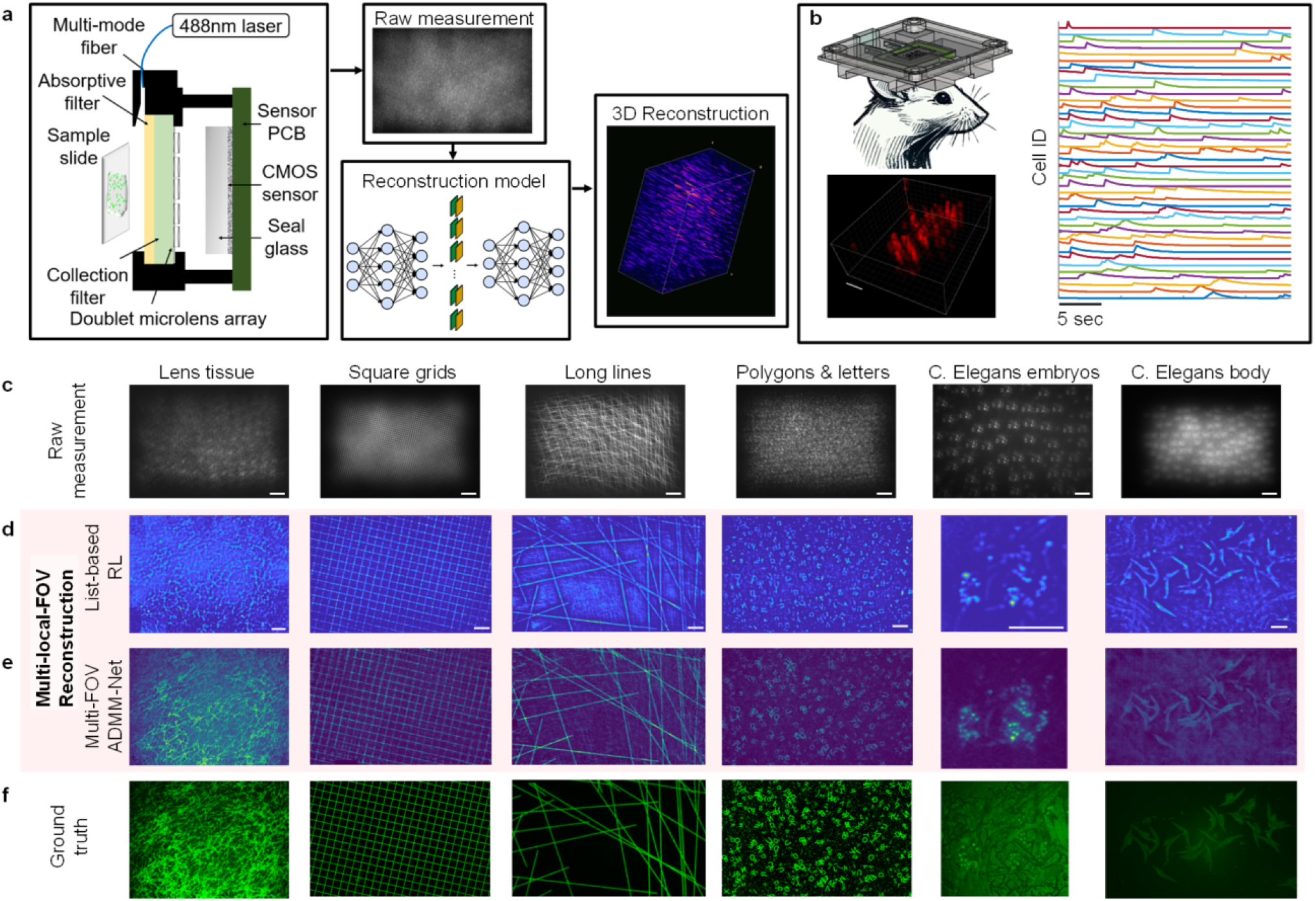
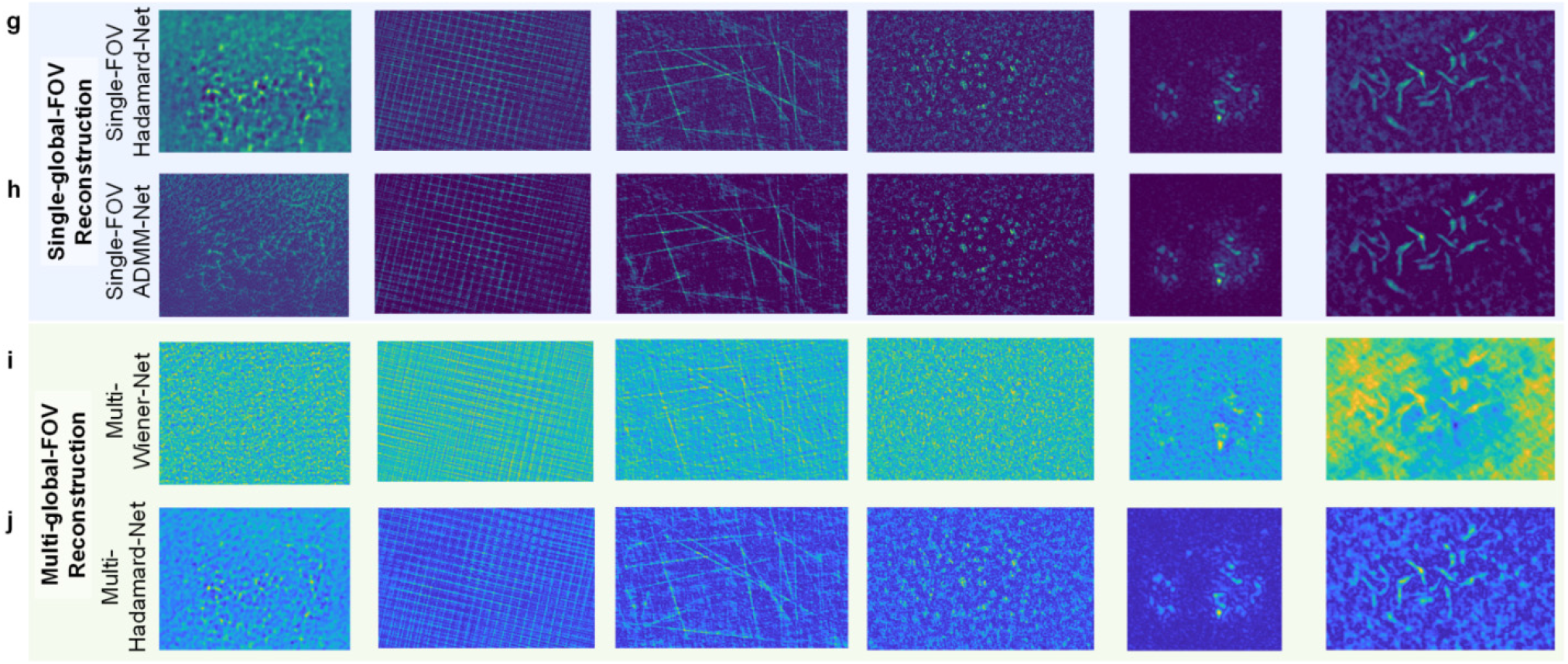
Overview of DeepLeMiN. (a) Schematic of DeepLeMiN. The raw measurement from the miniaturized lensless microscope is sent to a deep neural network, which reconstructs the object in 3D. (b) Application of DeepLeMiN in neural activity imaging in mouse brain in vivo. Left, a 3D volume in the visual cortex highlighting the reconstructed active neurons (shown as maximum projection of temporal activity); Right, the normalized extracted temporal activity of representative neurons. (c-j) Application of DeepLeMiN in fluorescence imaging. From left to right column, the samples are lens tissue, square grid patterns (12 µm line width), long line patterns (20 µm line width), polygons and letters, C. elegans embryos and C. elegans body. (c) Raw measurements by DeepLeMiN. (d) Reconstruction using a list-based RL algorithm. (e) Reconstruction using a multi-FOV ADMM-Net. (f) Ground truth obtained by a benchtop microscope. (g) Reconstruction using a single-FOV Hadamard-Net. (h) Reconstruction using a single-FOV ADMM-Net. (i) Reconstruction using a multi-Wiener-Net. (j) Reconstruction using a multi-Hadamard-Net. Scale bar, (b) 300 µm. (c)-(d) 500 µm.

## Results

### Miniaturized lensless microscope with a doublet microlens array

DeepLeMiN (Fig. 2, Methods) features a thin layer of microlens array as the optical mask. Each microlens unit is a custom-designed doublet lens (Fig. 2e-f) that minimizes wavefront aberrations, yielding a sharp PSF across an extended FOV. Compared with singlet lens, our design could achieve a higher Strehl ratio within the designed FOV while reduces the background outside its FOV (Fig. 2g). Each doublet lens has a diameter of 300 µm and a focal length of 1.17 mm, resulting in a numerical aperture (NA) of 0.1. The optical mask, consisted of 108 doublet lenses distributed in a semi-random pattern within an area of 4×6 mm^2^, was fabricated using two-photon polymerization^27^ on a fluorescence emission filter. The interspaces between doublet units were coated with aluminum to reduce the background light that is not modulated by the lenslets. We set the distance between the object and the microlens array as 4.9∼6.9 mm, and the distance between the microlens array and the image plane as 1.8 mm, resulting in a system magnification of 0.26-0.39. The airy radius of the doublet lens’ PSF was simulated to be 3.65 µm, corresponding to a lateral resolution ∼9.5 µm when considering the system magnification. We defined the FOV of each doublet to be ∼1 mm, where the peak intensity of the PSF drops to ∼0.8 of the value of the on-axis PSF, in good agreement with the simulation results of the design (Fig. 2g). This result confirmed that the PSF is spatially sparse with each lobe being individually constrained. As described later, this is a vital attribute that significantly decreases the computational demand for our reconstruction algorithm, setting our approach apart from others.

**Figure 2.**
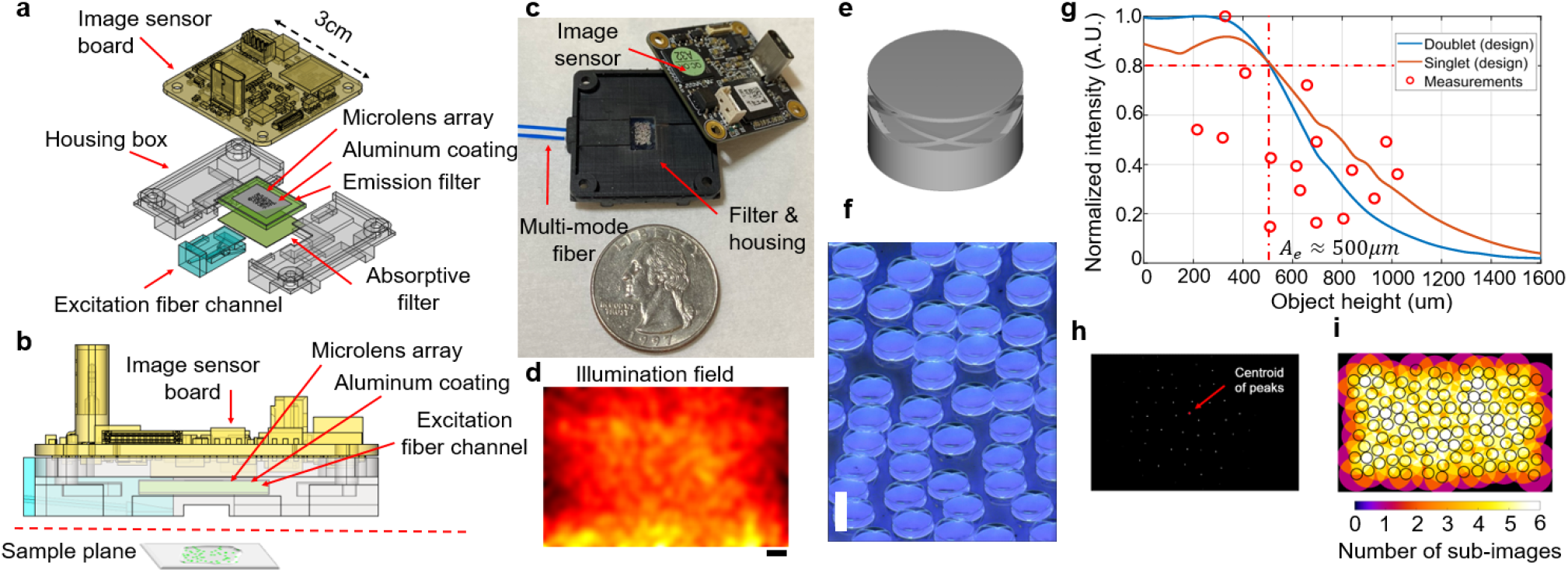
Assembly of DeepLeMiN and the microlens array. (a) Exploded-view of DeepLeMiN, illustrating the 3D printed housing, stack of optical filters with microlens array fabricated on top, and CMOS camera on the PCB board. (b) Side-view of DeepLeMiN. (c) Assembled device illustrating the imaging window with the microlens array. (d) Illumination intensity distribution across the field of view at sample plane with two fiber illumination channels, simulated by ray-tracing followed by a convolution with a Gaussian kernel with a size of 100×100 µm^2^. The raw results had discontinuity artifacts due to the limited number of rays, which could be suppressed by the convolution. (e) Structure of a doublet lets unit. (f) Fabricated microlens array observed from an optical microscope. (g) Normalized peak intensity of the PSF from a point source. Blue/red trace, simulated result from a doublet lens unit/singlet lens unit optimized for imaging quality within 500 µm object height. Dot, experimental measurement of the doublet lens unit from a point source in 4 µm diameter. The effective imaging area of the lens unit is defined when the peak intensity of the PSF drops below 80% of the maximum value. (h) Experimentally measured image of a point source in 4 µm diameter. (i) Number of sub-images obtained from the microlens array at each object location, assuming the effective field of view of each lens unit is 500 µm in radius. The microlens units are marked with black circles. Scale bar, (d) 500 µm, (f) 300 µm.

The microlens array on the emission filter is stacked with an absorptive color filter and assembled in proximity with a board-level back-illuminated CMOS camera sensor (5.5×7.4 mm^2^, 3000x4000 pixels) (Fig. 2a-c). An optional multimode fiber, bifurcating into two ports, can be inserted to the assembly to deliver the excitation light from an LED to the sample. The angle and position of the two fiber ports were optimized so the illumination is uniform across the FOV of 4×6 mm^2^ (Fig. 2d). The fluorescence signals pass through the filter stack and are captured by the doublet microlens array onto the camera sensor. The entire assembly is ∼30×30×6.5 mm^3^ in size with a weight of 7.5 grams (or 3 grams if excluding the camera).

### Highly efficient reconstruction algorithms

A major challenge in lensless microscopy is to reconstruct the high-quality 3D object while minimizing the computational resource and time, from a 2D image where the weak fluorescence signals are overshadowed by a strong background created by the overlap of the sub-images from different lenslets. There are two major classes of reconstruction algorithms: model-based iterative optimization methods such as those based on Richardson-Lucy or ADMM^7,8,13,24^, and deep learning methods utilizing neural network^17,25,26^. The former typically delivers higher and more consistent reconstruction quality but at the cost of substantial computational resource and time, along with the need for an accurate characterization of the PSF. Deep learning methods, on the other hand, once trained, can have a fast processing speed, though they often lack generalizability to data types beyond their training set and are constrained by the available GPU memory.

We develop multi-FOV ADMM-Net, a multi-stage and physics-informed deep neural network based on unrolled ADMM frameworks^22,23^ (Fig. 3, Supplementary Fig. S1-2, Methods). This network not only synthesizes the robust and high-quality reconstruction capabilities of ADMM-based iterative optimization with the high processing speed in neural networks, but crucially, significantly reduces the required computational resources, facilitating the reconstruction of large 3D data volume with a large voxel count from a 2D image with high pixel count.

**Figure 3.**
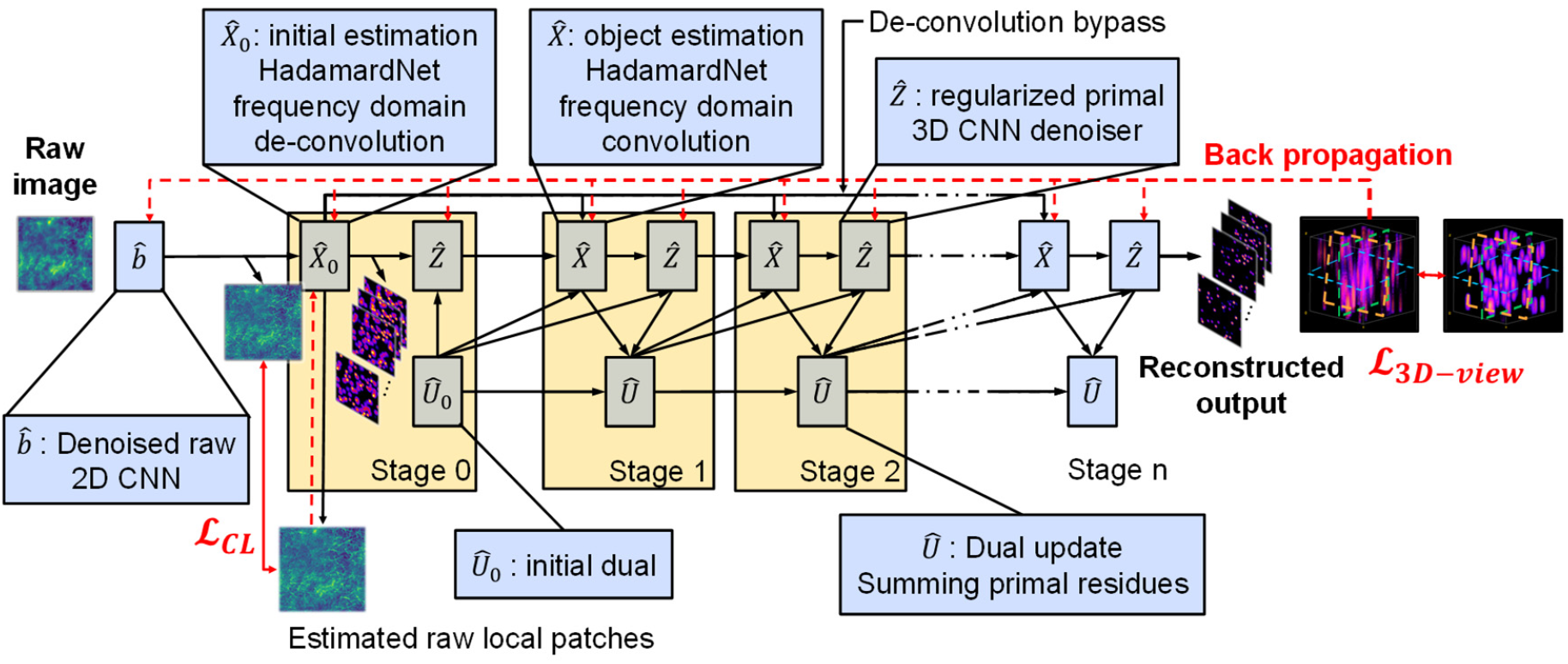
Architecture of ADMM-Net, illustrated as single-FOV to reconstruct a single 3D volume. The raw measurement is preprocessed by a 2D convolutional neural network, which could denoise and suppress the background of the raw image. The preprocessed raw image is then sent to the ADMM-Net, which contains multiple stages. Each stage contains a deconvolution module (stage 0) or convolution module (subsequent stages) to update the reconstructed image 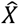, a CNN denoiser to update the regularized primal 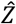, and a mathematical layer to calculate the dual variable / Lagrange multiplier *Û* .*ℒ*_*CL*_ is a closed-loop loss function defined as the structural similarity index (SSIM) between the estimated image based on the reconstruction and the denoised raw measurement. *ℒ*_3*D-view*_ is a loss function defined as the sum SSIM between the projected xy, yz, and xz view from the reconstructed volume and the corresponding ground truth.

The multi-FOV ADMM-Net starts with a pre-processing step for background suppression, utilizing either a U-Net or a Laplacian of Gaussian (LoG) filter, followed by a multi-head physics-aware neural network to initiate the reconstruction results. Each head of the network learns a local PSF and reconstructs the object within a localized FOV at a specific depth (Supplementary Fig. S3), through a Hadamard-Net in Fourier domain that we recently developed^16^. The Hadamard-Net, grounded in the underlying deconvolution principle of transpose convolution, learns the deconvolution process without requiring PSF calibration, making it highly efficient to create an initial reconstruction^16^. The preliminary reconstruction results are then refined through multiple ADMM-Nets (Supplementary Fig. S4), which has two distinct features. Firstly, each ADMM-Net reconstructs a 3D local FOV, offering unique advantages for microscopy lacking a global PSF (Supplementary Fig. S5). This allows us to model the image formation process in each local FOV through a unique PSF. Each ADMM-Net then incorporates this image formation into the learnable the network parameters through the local data patch (Supplementary Fig. S3). The local reconstructions are then fused together into an entire 3D volume and compared with the ground truth, from where a loss function is calculated and used to update the individual ADMM-Nets. Such a process cohesively integrated the individual FOV together. Secondly, each of our ADMM-Net has a significantly reduced demand of computational resources than the traditional ADMM-Net^22,23^. ADMM-Net itself has a relatively complex structure, which is composed of multiple iterative stages, including object updates through gradient calculation, regularization and denoising (through a convolutional neural network in our case), and dual update to account for primal residues (Methods). Typically, the computational challenge lies in the large memory needed in the object update stage which involves a multiplication of a linearized vector of the object and the inverse of a matrix (*A*^*T*^*A*+ *ρI*), where *A* is the system matrix describing how each object voxel is mapped to the image plane, *A*^*T*^ is the transpose of *A, I* is the indentity matrix at the same size of *A*, and *ρ* is the penalty parameter, a learnable scalar in the ADMM-Net. The computational complexity is *O*(*N*^2^), where *N* is the number of voxels in the object. For a large scale image data, for example in our case, *N*∼10^6^ - 10^7^. The multiplication would thus require ∼30 TB RAM in GPU, which is enormous. By exploiting the sharp and sparse PSF patterns, we can approximate (*A*^*T*^*A*+ *ρI*)^-1^ as an identity matrix. This allows us to transform this large matrix multiplication into the sum of multiple 2D convolution between the 2D object at each depth and a kernel which could be initialized as a 2D matrix with a single non-zero entry in the central element. This could be further simplified to Hadamard multiplication in Fourier domain (Methods). We can thus significantly reduce the computational complexity to *O*(*NlogN*), with a RAM requirement of only ∼ 24 GB. Additionally, our algorithm does not require any calibration of the PSF in experiment, which would be a laborious work in ADMM^13^ or typical deep neural networks ^17,23,25,26^.

We trained our neural network in experiment using custom-designed fluorescent samples, with ground truth images captured by a benchtop 2× microscope (Methods). This approach, distinct from conventional methods that rely on simulated data^17,26^, eliminates the need for extensive and precise calibration of the PSF. Furthermore, our method could better capture the image formation process which may not be fully modelled in simulation. Once trained, our network is capable of reconstructing a volume of 6×4×0.6 mm^3^ with ∼832×1248×13 voxels at 1 Hz (24 GB GPU RAM) or 3 Hz (80 GB GPU RAM).

To benchmark the performance of our neural network, we also developed a list-based Richardson-Lucy (RL) algorithm as an alternate method to reconstruct the objects (Supplementary Fig. S6, Methods). RL algorithm is a classical deconvolution method, but in 3D lensless microscopy, it would require too much memory to store the system matrix. Leveraging the sparse PSF, we turn the system matrix into individual lists, which contain the amount of camera pixels (<50) that each object voxel projects to. Effectively, this reduces the complexity of the system matrix from *O*(*N*^2^) to *O*(50*N*), thereby conserving memory and enabling the algorithm to operate on a standard laptop. Compared to the multi-FOV ADMM-Net, the list-based RL algorithm runs slower [12 second per iteration for 1.1×10^7^ voxels (16 planes)] and the reconstruction quality could be worse due to a lack of regularization. Nonetheless, the list-based RL algorithm is suitable for system lacking a global PSF, and has a generally robust reconstruction performance, which could serve as a baseline to evaluate the performance of the Multi-FOV ADMM-Net.

### Image resolution

We characterized the image resolution using both a single point source and a resolution target (Supplementary Fig. S7-8). The lateral resolution of our lensless imager is ∼10 µm, verified by imaging a 4 µm point source, and the USAF resolution target (group 5, element 6, ∼9 µm line width), which is consistent with our designed value. The axial resolution is ∼60 µm. These results have the potential for improvement by enhancing the uniformity in the fabrication of the microlens units.

### Imaging of phantom with dense fluorescence features

We validated DeepLeMiN and the reconstruction algorithms on fluorescence samples such as lens tissue stained with fluorescent spray, and customized-fabricated masks containing letter shape features, mesh grids, or randomly distributed lines (Fig. 1c-f). The lens tissue posed a significant challenge for reconstruction due to its dense fluorescence features, leading to a strong background in the raw image. Before the actual reconstruction, we used a U-net to suppress the background (Methods). For other samples, we used LoG filters to highlight the edges, which could facilitate the subsequent reconstruction. While both the multi-FOV ADMM-Net and list-based RL algorithm successfully reconstructed the object, the multi-FOV ADMM-Net demonstrated superior performance across various evaluation metrics (Supplementary Fig. S9), including both spatial aspects [such as peak-SNR (PSNR) and structural similarity index (SSIM)] and frequency domain measures (Methods).

We further compared the performance of the Multi-FOV ADMM-Net with networks that reconstruct the image with a global PSF or FOV (Fig. 1g-j), such as a single-FOV ADMM-Net (Fig. 1h) or a single-FOV Hadamard-Net^16^ (Fig. 1g), and a multi-Wiener-Net^26^ (Fig. 1i) or a multi-Hadamard-Net^16,28^ (Fig. 1j). The single-FOV ADMM-Net or Hadamard-Net, limited to learning a single PSF for the entire FOV, could only achieve a satisfactory reconstruction within the central FOV. The multi-Wiener-Net or multi-Hadamard-Net (with kernels randomly initialized as in the other algorithms in comparison), designed to learn multiple global PSFs and to merge multiple reconstructions of the entire FOV, are not exactly suitable for our imaging system and resulted in strong background artifacts. The Multi-FOV ADMM-Net, on the other hand, resulted in reconstructions with a more balanced quality and intensity and a reduced background across the entire FOV, particularly in the samples with dense and complex features such as the lens tissue.

### Imaging of fluorescent beads embedded in scattering medium over a large 3D volume

We tested the 3D imaging capability of DeepLeMiN on phantoms containing randomly distributed fluorescent beads (12 µm diameter) in clear media (Fig. 4a-d). We first used a benchtop microscope with a 2× or 10× objective lens to characterize the sample (Fig. 4a). The former has a large FOV but poor 3D resolving power, whereas the latter can resolve multiple axial depths by changing the focus but has a smaller FOV. DeepLeMiN combines the strength of both objective lenses, and could resolve 3D features over a large volume (6×4×0.6 mm^3^, 13 planes, reconstructed by either Multi-FOV ADMM-Net or list-based RL). The z-projection aligns well with that obtained from the benchtop 2× microscope, and the axial distribution of the beads agrees with that from the 10× microscope (Fig. 4b-c). We analyzed the histogram of the axial full-width-at-half-maximum of the reconstructed beads, and confirmed that the minimum axial resolution was ∼50 µm (Fig. 4d).

**Figure 4.**
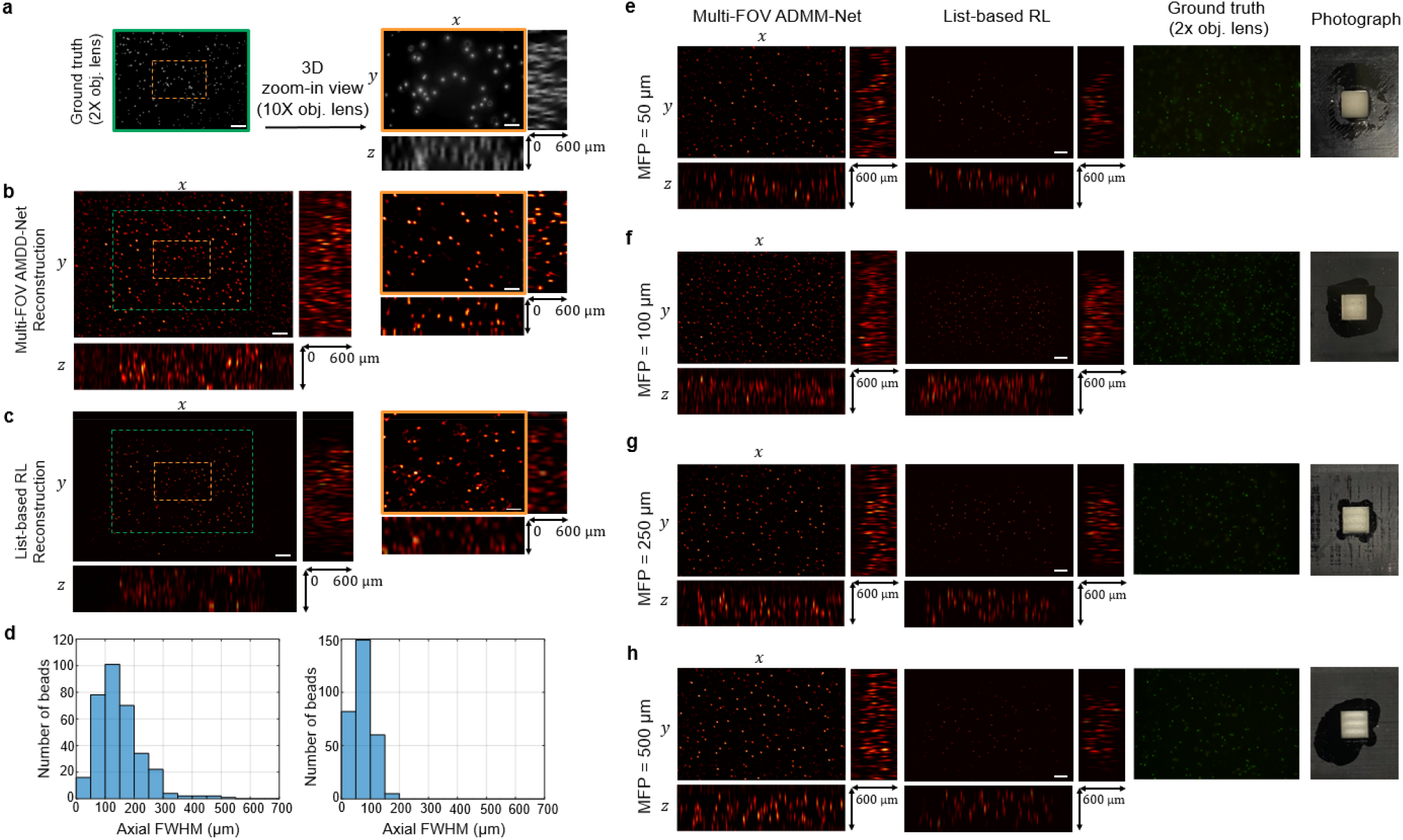
3D reconstruction of fluorescent beads distributed in a 3D volume. (a) Fluorescent beads phantom (fluorescent beads in 12 µm diameter distributed in optical clear polymer) imaged by a benchtop microscope with 2× objective lens (left, XY view) and 10× objective lens (right, XY, YZ, XZ view, 13 axial planes, each separated by 50 µm). The image from the 10× objective lens (right) is a zoom-in view of the region inside the orange dashed box in the image from the 2× objective lens (left). (b) 3D reconstruction results within 600 µm depth range through a multi-FOV ADMM-Net, in XY, XZ, YZ views. The green/orange dashed box in the XY view (left) corresponds to the field of view of the image from the 2×/10× objective lens respectively in (a). Right, zoom-in view of the region inside the orange dashed box in the left image. (c) Same as (b), but with list-based RL algorithm. (d) Histogram of the axial FWHM of individual beads reconstructed from the multi-FOV ADMM-Net (left) and list-based RL algorithm (right). (e-f) 3D reconstruction of fluorescent beads distributed in a 3D scattering volume with a mean-free-path (MFP) of (e) 50 µm, (f) 100 µm, (g) 250 µm, and (h) 500 µm. The optical clear polymer is mixed with fluorescent beads in 12 µm diameter and non-fluorescent beads in 1.18 µm in diameter, whose concentration could be used to control the mean-free-path (MFP). All reconstructions are within 4.2 mm×5.8 mm×600 µm volume range. Left/middle, reconstruction through multi-FOV ADMM-Net/list-based RL algorithm, in XY, XZ, YZ views. Right, reference image captured by a benchtop microscope with a 2× objective lens, and the photograph showing the scattering phantom slide on top of a resolution target. Scale bar, (a)-(c) left, 500 µm; right, 200 µm. (e)-(h), 500 µm.

In subsequent experiments, we imaged 3D scattering phantoms where the 12 µm diameter fluorescent beads were mixed with 1.18 µm diameter non-fluorescent beads, which act as scatterers (Fig. 4e-h). As the scattered light increases the imaging background, we first removed the background from the raw image by a LoG filter, before processing them with the multi-FOV ADMM-Net or list-based RL algorithm (Methods). Both reconstruction methods could pick up the ballistic light, and reconstruct the 3D volume across scattering lengths of 50∼500 µm. The results are in a good agreement with the images captured by the benchtop microscope, demonstrating the robustness of reconstruction algorithms.

### Imaging of C. Elegans and Hydra

Imaging the biological specimens presents challenges for lensless imager due to the typically weak fluorescence signals and the presence of background fluorescence over the regions of interest. Our reconstruction algorithm, equipped with a background removal module and multi-FOV ADMM-Net, is particularly tailored to tackle this challenge. The algorithm iteratively learns the image formation and progressively enhances the reconstructions. We validated the reconstruction capability by imaging C. elegans (Fig. 1) and the 3D motion of live hydra (Fig. 5), where the embryos of the C. elegans and the endodermal or ectodermal epithelial cells of the hydra were labeled with green fluorescent protein (GFP). In C. elegans, our system was capable of distinguishing embryos as small as 13 µm. In hydra, it successfully captured its tentacles with thicknesses down to ∼55 µm. Moreover, we recorded the 3D motion of the hydra, across a FOV ∼6×4×2 mm^3^ at 10 Hz, reconstructed using list-based RL algorithm. While traditional benchtop microscopes are commonly used to study the morphology of hydra without axial resolution, our lensless microscope offers a distinct advantage of high-resolution 3D imaging over large volumes.

**Figure 5.**
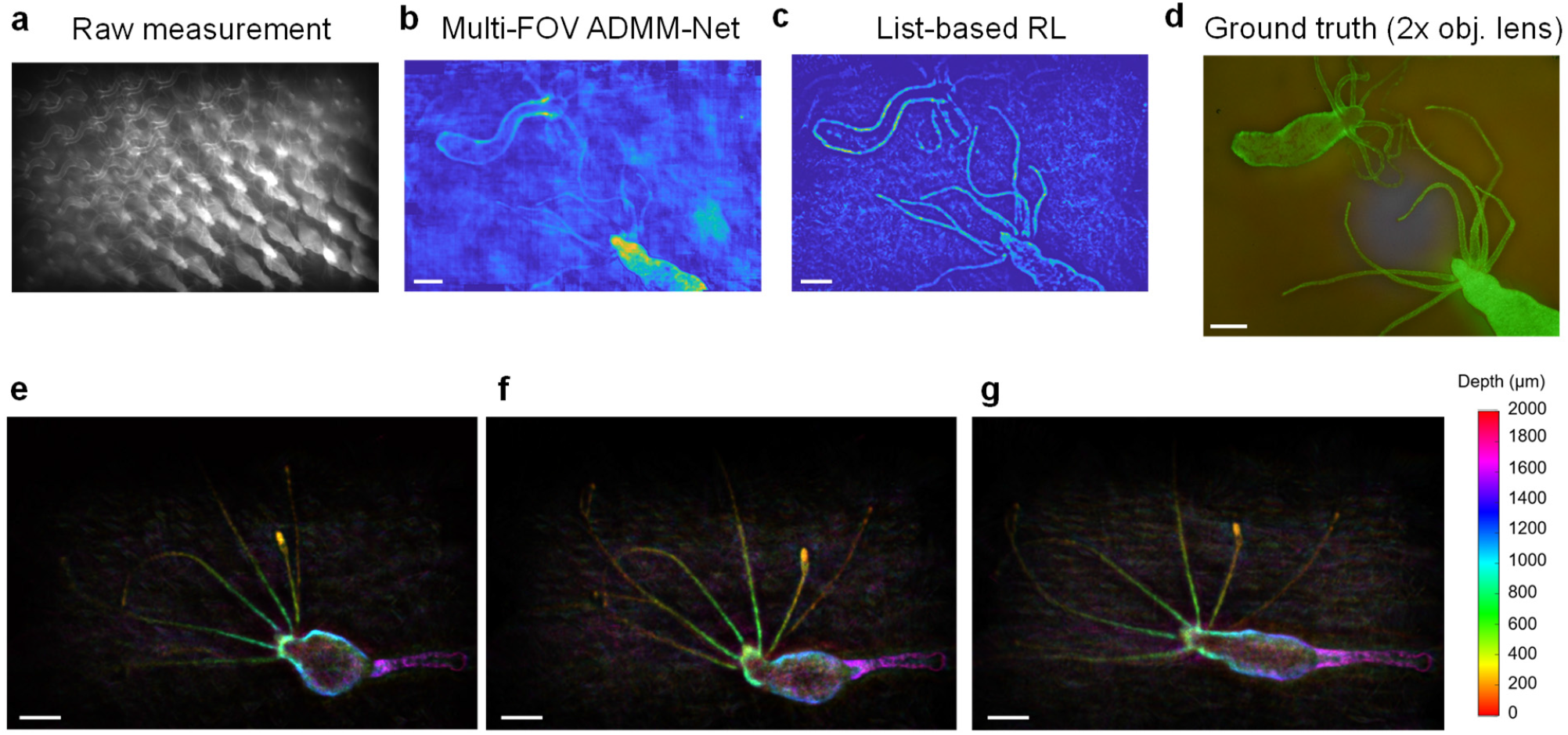
In vivo 3D imaging of hydra labeled by green fluorescent protein (GFP). (a-d) Image and reconstruction of two hydras, with the top one having ectodermal cells labeled with GFP, and the bottom one having endodermal cells labeled with GFP. (a) A raw image frame from DeepLeMiN. (b) 2D reconstruction of (a) by the list-based RL algorithm. (c) 2D reconstruction of (a) by the multi-FOV ADMM-Net. (d) Reference image captured by a benchtop microscope with 2× objective lens, at a slightly different time from that captured by DeepLeMiN, as the two images cannot be captured at the same time. (e-g) 3D reconstruction of a hydra at three different frames over 2 mm axial range, by list-based RL algorithm. The color bar indicates the reconstruction depth. Scale bar: (b)-(g) 500 µm.

### Imaging of the visual cortex of awake mouse

One key application of DeepLeMiN is to record neuronal activity over a large FOV in awake mouse. We conducted in vivo experiments to monitor the spontaneous activity of layer 2/3 in primary visual cortex (V1) in head-fixed awake mouse transfected with calcium indicators GCaMP6f^29^ (Fig. 1b, Fig. 6). A significant hurdle encountered in imaging scattering brain tissues was the pronounced background in the raw recordings, which prevented imaging at the single-cell level in previous studies^15,19^. To address this, we preprocessed the raw recording by suppressing the pixels exhibiting small temporal variation (Methods). Subsequently, the data was processed through the Multi-FOV ADMM-Net. We successfully extracted ∼150 regions of interest with cellular resolution, over a 3D volume spanning 1.5mm×2mm×300µm (limited by the spatial expression range of GCaMP6f and the size of craniotomy in our current animal preparation protocol). Using a constrained negative-matrix factorization algorithm (CNMF-E)^30^, we deconvoluted the activity of individual neurons. This represents, to the best of our knowledge, the first in-vivo calcium imaging at cellular resolution in mouse brain achieved with a lensless miniaturized microscope.

**Figure 6.**
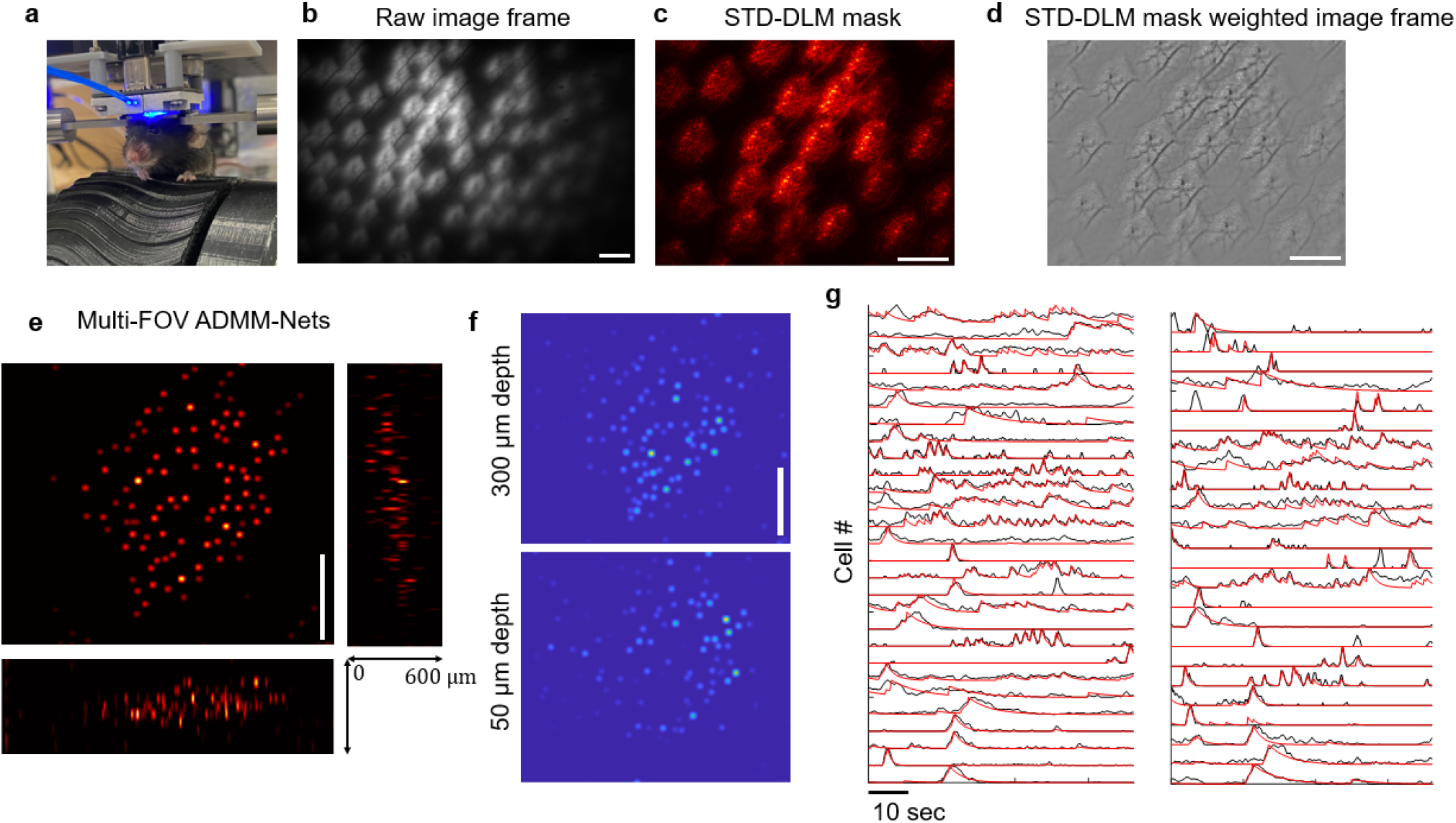
In vivo calcium imaging of neural activity in mouse visual cortex, transfected by GCaMP6f. (a) Experimental setup of the in-vivo calcium imaging. The mouse was head-fixed on a treadmill, with DeepLeMiN mounted on top of the headplate. The excitation light was delivered through the dual fiber channels. (b) A single raw image frame. (c) Time-series standard-derivation of difference-to-local-mean of the raw video, termed as STD-DLM mask. The STD-DLM mask highlights the pixels with strong temporal dynamics. (d) A single raw image frame of the raw video weighted by the STD-DLM mask. (e) 3D reconstruction of the image volume (1.5mm×2mm×300µm) by a multi-FOV ADMM-Net, in XY, YZ and XZ view, showing the time-series standard-derivation projection of the reconstructed video. The multi-FOV ADMM-Net took the STD-DLM-mask weighted frames as input. This spatial reconstruction volume was further processed by an iterative clustering algorithm^14^ to highlight individual neurons. (f) Two individual axial planes from (e). Top, 300 µm depth. Bottom, 50 µm depth. (g) Representative normalized temporal activity traces of the extracted neurons. Black, traces of the neurons from the reconstructed video. Red, deconvoluted traces through CNMF-E^30^. Scale bar: (b)-(f) 500 µm.

## Discussion

In summary, we developed DeepLeMiN, a miniaturized lensless microscope with a custom-designed doublet microlens array. Our innovative multi-FOV ADMM-Net algorithm is capable of reconstructing local FOVs and synthesize them to a larger FOV, demonstrating the scalability of the imaging FOV. We showed that the multi-FOV ADMM-Net was highly efficient in computation, and had superior reconstruction capabilities over previous methods. We tested DeepLeMiN across a variety of samples, including those with dense and detailed features over expansive areas such as lens tissue, and biological specimens where the regions of interest exhibited weak fluorescence or were embedded in scattering tissues such as neurons in awake mice. Additionally, we introduced a list-based RL algorithm, and successfully reconstructed the 3D motion of a hydra over a 2 mm depth of field.

One key contribution of our work is the development of the multi-FOV ADMM-Net, designed for imaging systems that do not have a global PSF. Typical deep learning algorithms for lensless imagers adopt a single PSF kernel to reconstruct images. While certain algorithms like multi-Wiener-Net^26^ utilize multiple PSF kernels to improve the reconstruction in spatial variant systems, they apply each PSF kernel globally. The reconstructed image through each PSF has high quality only within a specific area. These areas are then combined to form the final image. Such a global reconstruction approach is inefficient if the PSF only encodes a point source to a nearby region on the image sensor. Our method innovatively partitions the raw data into localized patches for individual reconstruction, and subsequently merges them into a large FOV output. This strategy of focusing on smaller, local patches significantly improves the quality and efficiency of reconstruction, making our algorithm particularly suited for our miniaturized microscope with a scalable FOV.

Our method shares similarity with prior strategies that divide the FOV into localized images for each lens unit, and employ the light field imaging technique for 3D volume reconstruction through refocusing^17^. However, our approach distinguishes itself by performing deconvolution on each localized FOV, rather than merely demixing them at the image sensor plane. This crucial advancement enables our system to tackle the more complex imaging scenarios where the local images of each lens unit significantly overlap, such as imaging samples with dense features spanning a large area.

Our work also presents a significant advancement in optimizing the computational efficiency of the ADMM-Net, which leverages the unique attributes of our PSF. Traditionally, ADMM-Net demands substantial memory resources as it is unrolled into multiple stage, each involving a large-scale matrix multiplication. In our case, the sparse and spatially discrete nature of the PSF from our doublet microlens array allows an innovative approximation of *A*^*T*^*A* as an identity matrix. This faciliates the conversion of the large matrix multiplication into 2D convolutions, which is then calculated by more manageable Hadamard multiplications in Fourier domain. Such a strategy reduces the computation complexity by orders of magnitude. Furthermore, this approach eliminates the need of a specific initialization of system matrix *A*. In conventional ADMM or deep neural network such as the multi-Wiener-Net, *A* is either fixed or requires careful initialization through an extensive PSF calibration across the entire object domain, which is time consuming.

Our miniaturized lensless microscope, constructed with a doublet microlens array and complemented by a highly efficient multi-FOV ADMM-Net reconstruction algorithm, allows high resolution 3D microscopic imaging over a large FOV. The strategy that decomposes the entire view into local patches for individual construction before stitching them together, establishes a new platform for microscopy. This approach fundamentally allows for the FOV to be scaled up unlimitedly. Future work could explore integrating the microlens design within a differentiable neural network framework to jointly optimize the imaging optics and reconstruction, and demonstrating imaging in freely-behaving animals.

## Acknowledgement

This work is partially supported by National Eye Institute (R21EY029472), Burroughs Wellcome Fund (Career Award at the Scientific Interface 1015761), and National Science Foundation (CAREER 1847141). We acknowledge Dr. Celina Juliano and Dr. Ben Dillon Cox for preparing and sharing the hydra samples, Dr. Francis McNally and Elizabeth Beath for preparing and sharing the C. Elegans samples, Dr. Lin Tian for providing the surgical setup and tools for mice surgery, and Center for Nano-MicroManufacturing at UC Davis for the support in fabricating the doublet microlens array.

## Methods

### Construction of the miniaturized lensless microscope

The doublet microlens array was fabricated on a fluorescent long-pass filter (ET525/50m, Chroma), with the interspaces between the lenslets coated with aluminum to reduce the background light that is not modulated by the lenslets. The fabrication process was separated into two steps. First, photolithography was performed on the filter glass to define the interspaces between the lenslets. An 80 nm thick aluminum layer was then deposited to the filter through electron beam evaporation, followed by liftoff (Supplementary Fig. S10). Such an aluminum layer could reflect >99% of the fluorescent light and excitation light, supported by Finite-Difference Time-Domain (FDTD) simulation (Supplementary Fig. S11). Second, the doublet microlens array was fabricated by two-photon polymerization process^27^ (Lumenworkx). To support the overhang microlens component on top, we incorporated dual helix structures to connect the edges of two lens components, which could maximize the support strength while allowing sufficient access for developers to remove the photoresist in between the two lens components. The bottom piece of the array was attached directly onto the glass surface free of aluminum deposition. The dual helix support structures and the top components of the lens units were then added atop the bottom components.

The doublet microlens array was then stacked on top of an absorptive filter (Yellow 12 Kodak Wratten color filter, 14×11.3×0.1 mm^3^, Edmund Optics). The entire stack and a board-level back-illuminated CMOS image sensor (IMX226, The Imaging Source) were then assembled in a 3D printed housing (AlSi10Mg, Protolabs). Two fiber ports (Ø400 µm, Thorlabs) could be inserted to the assembly to for sample illumination.

### Architecture of ADMM-Net (single FOV reconstruction)

The ADMM-Net is an unrolled alternating direction method of multipliers (ADMM) with learnable parameters, aiming to recover the 3D object *X* from a measured image *b* of an optical imaging system with a forward model of *b* = *AX* + *n*, where *A* is the system matrix and *n* is the noise. Such a reconstruction problem can be generally expressed as an optimization problem:

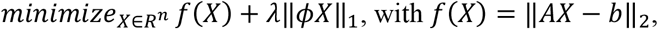

where *λ*‖*ϕX*‖_1_ is a regularization term, *ϕ* is an operator that transforms *X* into a sparsity representation, and *λ* is a hyper-parameter.

The ADMM algorithm formulates this optimization problem as

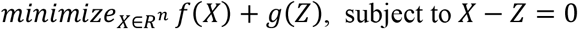

where *Z* is the regularized variable of *X*, and *g*(*Z*) is the regularization term of *Z* (and thus *X*).

The augmented Lagrangian in its scaled form is

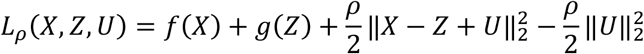

where *U* is the dual variable or Lagrange multiplier, and *ρ* is the penalty parameter.

The ADMM algorithm then iteratively updates *X, Z* and *U* through the following steps (in *k*^*th*^ iteration, *k* ≧ 0)^31^:

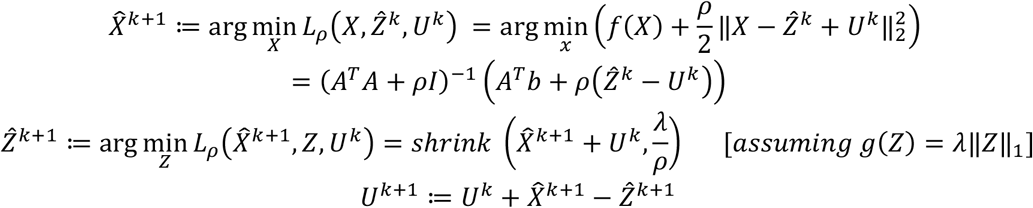

where *shrink*(*X, k*) = sign(*X*)max(|*X*| - *κ*, 0) is a soft thresholding operator.

The ADMM-Net^22^ contains multiple stages, with each stage representing an iteration of the ADMM algorithm. It allows the parameters of the ADMM algorithm, as well as the regularization term, to be learnt from the data, offering improved performance, flexibility, and efficiency. In conventional ADMM-Net for image reconstruction, both *X* and *Z* are vectorized version of the 2D/3D image, and *A* is a 2D matrix with a large dimension. Storing *A* and operating on *A* require a large amount of memory and are computationally expensive. Furthermore, *A* needs to be initialized by the PSF. For the spatial variant system, this is a laborious process in experiment.

Our realization of ADMM-Net does not require vectorizing *X* and *Z*, or storing matrix *A*, and can turn the large matrix multiplication into convolution operations which could be performed in two-dimensional frequency domain. This significantly reduces the computation complexity from *O*(*N*^2^) to *O*(*NlogN*) [assuming *N* is the number of voxels in the object, which is on the same order as the number of pixels in the recorded image], and thus the required memory and computational resources.

We developed two types of ADMM-Net: type I is more standard and close in format to the conventional optimization, and type II demands less memory (Fig. 3, Supplementary Fig. S1-S2). We first describe their structures for 2D sample reconstruction, and then 3D sample reconstruction.

In Type I ADMM-Net, the update of *X, Z* and *U* in each stage can be described as:

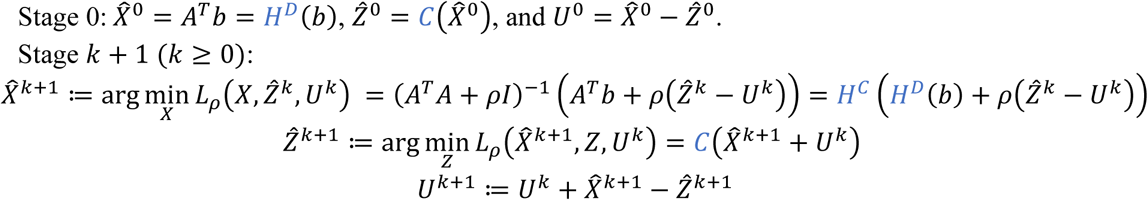

where *H*^*C*^, *H*^*D*^, *C* and *ρ* represents trainable modules or parameters for each stage.

*H*^*D*^ represents the deconvolution Hadamard layer (Hadamard-Net)^16^, which operates in frequency domain and deconvolutes *b* into *X*, both of them in a format of 2D matrix (Supplementary Fig. S1). It aims to recover *A*^*T*^*b*. As the PSF in our system is sparse and discrete, *A*^*T*^ ≈ *A*^-1^, so *H*^*D*^(*b*) = *A*^*T*^*b* ≈ *A*^-1^*b*. The design of *H*^*D*^closely resembles the deconvolution process in Fourier domain for optical imaging systems. It first transforms *b* into 2D Fourier domain, and then performs an element-wise multiplication (i.e. Hadamard multiplication) between the 2D Fourier transform of *b* and 2D learnable kernels, respectively for both real components and imaginary components. It then transforms the result back to the spatial domain, which is a 2D matrix. The weights of the kernel are randomly initialized.

*H*^*C*^ represents the convolution Hadamard layer (Supplementary Fig. S1). As our PSF is sparse and spatially discrete, *A*^*T*^*A* can be approximated as a 2D identity matrix, so does (*A*^*T*^*A*+ *ρI*)^-1^. 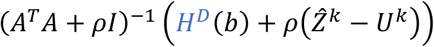 can thus be treated as a 2D convolution between a kernel associated with (*A*^*T*^*A*+ *ρI*)^-1^ and the 2D matrix 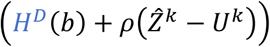. As (*A*^*T*^*A*+ *ρI*)^-1^ could be approximated as a 2D identity matrix, the kernel could be initialized as a 2D matrix with a single non-zero entry in the central element. This convolution could then be calculated in frequency domain through element-wise multiplication (i.e. Hadamard multiplication) where the kernel (in frequency domain) could be learnt. The weights of this kernel in frequency domain could be initialized as 1 for all the elements.

*ρ* is a learnable parameter initialized as a random scalar between 0 and 1. It is used to weight the regularized solution when being added to *H*^*D*^(*b*).

*C* represents a 2D convolutional neural network, serving as a learnable regularizer model (Supplementary Fig. S1).

In Type II ADMM-Net, the overall network architecture is the same as type I but requires less memory. As the deconvolution network *H*^*D*^ is designed based on the physics of the optical imaging system, we can safely replace *H*^*D*^ in all the stages by 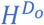 from stage 0. In other words, all *H*^*D*^ shares the same parameters as 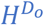. This reduces the required memory for our network. The update of 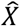 can be described as below:

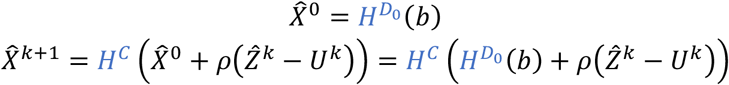

The updates of 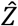 and *U* remain the same as Type I ADMM-Net.

In this paper, we adopt Type II ADMM-Net to save the computation resource. There were a total of six stages in the ADMM-Net.

For 3D samples, we build an individual 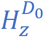 to reconstruct the object at individual depth *z*. Its output is then cropped and rescaled based on the magnification of each depth, resulting in 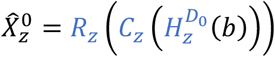, where *C*_*z*_ and *R*_*z*_ correspond to learnable center crop layers and resizing layers respectively for depth at *z*. This keeps the reconstruction of all axial depth in the same magnification factor, avoiding the lateral position shift between depths. At stage *k*, 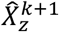 is updated by individual 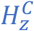, following the same procedure as the 2D reconstruction described above. Individual 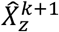 can be concatenated into a 3D matrix 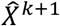. The update of 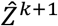 and *U*^*k*+1^ follow the same procedure as the 2D reconstruction, except that 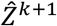 and *U*^*k*+1^ is now 3D matrix, and *C* is a 3D convolution neural network.

An extra step in 3D reconstruction is to calculate a predicted measured image 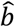, which will be used in calculating loss function, as described later. 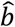 could be obtained through the learnt 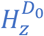 and 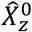. Based on 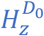, we could construct a forward model of the imaging system for the corresponding depth *z*. This forward model takes the complex conjugate of the weights learnt in 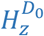, and multiple it with the Fourier transform of 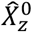 to obtained a predicted 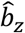 . The predicted 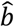 can then be obtained by

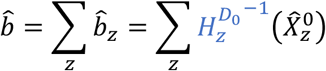

We built ADMM-Net stage by stage and trained it iteratively with an increasing number of stages. For 3D reconstruction, we pre-trained 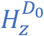 independently for each depth as 2D reconstructions, with a 2D binary cross entropy (BCE) loss between 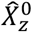 and ground truth *X*_*z*_ for each depth *z*. We then incorporated the 3D imaging data, and trained all 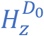 in stage 0 with a 2D binary cross entropy (BCE) loss between 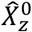 and ground truth *X*_*z*_ for each depth *z*, as well as a structural similarity index (SSIM) loss between 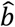 and the real measurement *b*. The latter loss function could increase the robustness of the network and its axial sectioning capability. We then added and trained *C* in stage 0 with a 3D BCE loss between 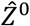 and 3D volume ground truth *X*, and again SSIM loss for 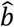. We then cascaded stage 1 and trained both stage 0 and 1 together with 3D BCE loss for 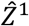 and SSIM loss for 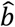. We then added subsequent stages and performed training with SSIM loss for 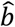 and a new 3D view loss for 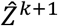 calculated in the last stage. The 3D view loss is defined as the sum SSIM loss between the projected xy, yz, and xz view and the corresponding ground truth. This loss function can enhance the axial sectioning capability. We chose to apply this loss function only after more stages were added to the models so that the model could focus to learn the baseline reconstruction, before enhance the axial resolution. For 2D reconstruction, only the 2D BCE loss function for 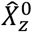 and SSIM loss 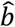 were used. As ADMM naturally converges to optimal relatively fast, in our realization we had a total of 6 stages in ADMM-Net to save the GPU memeory.

### Multi-ADMM-Net (multiple local FOVs reconstruction)

For 2D reconstruction, the entire FOV (4×6 mm^2^) was divided into 108 FOVs, each being 1.3×1.3 mm^2^ in size and centered at a single microlens unit (Supplementary Fig. S3). For 3D reconstruction, the entire volume (4×6×0.6 mm^3^) was divided into 6 sub-volumes (limited by GPU RAM), each being 2.02×2.02×0.6 mm^3^. While each FOV/sub-volume was reconstructed through a single ADMM-Net, the loss functions were globally calculated across the entire FOV/sub-volume, and *C* was shared in the same stage among all the ADMM-Nets. The output of individual FOV/sub-volume was then stitched together where the overlapped regions were averaged.

### Denoise module

A denoise module before the Multi-FOV ADMM-Net could be used to pre-process the raw images to suppress the noise/background and pick up useful features. This module is optional and could be realized by various algorithms, such as Lapacian-of-Gaussian (LOG), edge filtering, or a pre-trained convolutional neural network (CNN), depending on the exact images. For images containing dense features that are not obviously regulated in any domain, a flexible preprocessing module such as a UNet could enhance the overall reconstruction results of multi-FOV ADMM-Net (Supplementary Fig. S1). We first trained 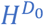 without this UNet. Using the noisy raw images as the input, 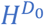 will learn to reconstruct the object with a rough estimation under the noise. Next, we connected the untrained UNet in front of the pre-trained 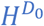, and trained the two modules together. Such a procedure avoids the scenario that the untrained 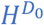 and UNet confuses each other and does not process the image correctly. After a few epochs training with two modules connected, the UNet can learn to extract useful features from the raw measurements even if there is no ground truth for the output of UNet, and 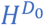 will adopt the output of UNet and enhance the reconstruction quality.

### Training dataset

We created training dataset by capturing images of a 2D fluorescent phantom samples using a bench top inverted microscope and DeepLeMiN. The phantom samples were features such as letters and polygons on a mask (prepared by spraying green fluorescent paint on a custom-designed photolithography mask) (Supplementary Fig. S2). For each image reconstruction depth, we accordingly set the distance between the miniaturized lensless microscope and the sample. We then performed two calibration steps to align the image from the benchtop microscope (ground truth) and that of DeepLeMiN (input of the reconstruction algorithms), using two calibration masks. The first calibration mask is a 4×6 mm^2^ rectangular mask with marks on the 4 corners. Using these marks, we could define the effective region on DeepLeMiN’s image which represents a 4×6 mm^2^ FOV. This effective region (2048×3072 pixels) then serves as the paired image of the ground truth. The second calibration mask contains dots separated by 600 µm. This could calibrate the magnification of DeepLeMiN. We also calculated the relative rotation angle between the images from the two microscopes and corrected such a rotation. After these two calibrations, we captured 15 images pairs for different phantom features (or the same features but different rotations) from both microscopes. This completed the acquisition of training data for one imaging depth. Then we displaced DeepLeMiN to change its distance from the sample, while keeping the sample and the inverted microscope in the same position, and repeated the steps above to acquire training data for another imaging depth.

Once all the training pairs of different imaging depths were obtained, we could synthesize 3D training pairs where the images of DeepLeMiN were a summation of the images of individual depth.

For each training pairs in 2D reconstruction, we divided the entire FOV into 108 local FOVs, each centered at a lens unit. Each local FOV is 1.3×1.3 mm^2^ in the raw measurement, and 1×1 mm^2^ in the output of ADMM-Net and ground truth. The larger size of the image in the raw measurement patch is to include all the sub-images of the microlens units within the local FOV for reconstruction algorithm.

For each training pairs in 3D reconstruction, we divided the entire volume (4×6×0.6 mm^3^) into 6 sub-volumes (limited by GPU RAM). Each local FOV is 2.6×2.6 mm^2^ in the raw measurement, and each sub-volume is 2.02×2.02×0.6 mm^3^ in the output of ADMM-Net and ground truth.

The 2D multi-FOV ADMM-Net model is trained on Tesla A10G (24GB RAM), and the 3D multi-FOV ADMM-Net model is trained with Tesla A100 (80GB RAM).

### List-based Richard-Lucy (RL) deconvolution

RL deconvolution is an iterative algorithm to reconstruct the object using the point spread function. Instead of using a full system matrix to store the mapping relationship between the object voxel and camera pixel, we use two sets of lists (Supplementary Fig. S6). The first sets of lists store the contributive object voxels for each camera pixel, whereas the second sets of lists store the contributive camera pixels for each object voxel (i.e. the PSF). This significantly reduces the required memory and computational time.

The two sets of lists could be obtained using geometrical optics (Supplementary Fig. S6). We set up a global coordinate across the object space and image space, with the lateral grid size being equal to the size of the pixel in the camera sensor. The nominal FOV is set to be 4×6 mm^2^, same as the size of the microlens array. We could expand the FOV by a ratio of *S*_*x*_ and *S*_*y*_ respectively in the *x* and *y* direction, as object points that are laterally outside the microlens array area could be imaged by some lens units at the boundary onto the camera sensor. In our setup, *S*_*x*_ and *S*_*y*_ was <20%.

The pixel size on our camera sensor was 1.85×1.85 µm^2^, and the magnification of the imaging system was 0.26-0.39. This translates to the lateral size of the voxel in the object space to be 4.7∼7.1 µm. We could thus down-sample the global coordinate by a factor of *D*_*S*_ in the lateral direction as the coordinate of the object space, while keeping the same FOV. In other words, within the global coordinate system, we picked one point out of every *D*_*S*_ x *D*_*S*_ points as the voxel to be reconstructed. Given the magnification and resolution, we set *D*_*S*_ to be 3.

For each object voxel, we used ray-tracing approach to find out the contributive camera pixels. We approximated the PSF of each lens unit to be a single point. As we have 108 microlens units, there will be a maximum of 108 pixels. However, as the object voxel is close to microlens array, only a portion of the lens units could effectively image a single object voxel (Fig. 2g-h, Supplementary Fig. S5). Correspondingly, pixels in the camera sensor that were laterally far away (*d*_*max*_ pixel counts away) from the object voxel of interest would not be considered. In our setup, *d*_*max*_ was set to be ∼800-1000 pixel units.

The pseudo code below illustrates the process to find the two sets of lists.

#### Algorithm 1: Establishing the two sets of lists (voxel-pixel mapping and pixel-voxel mapping)

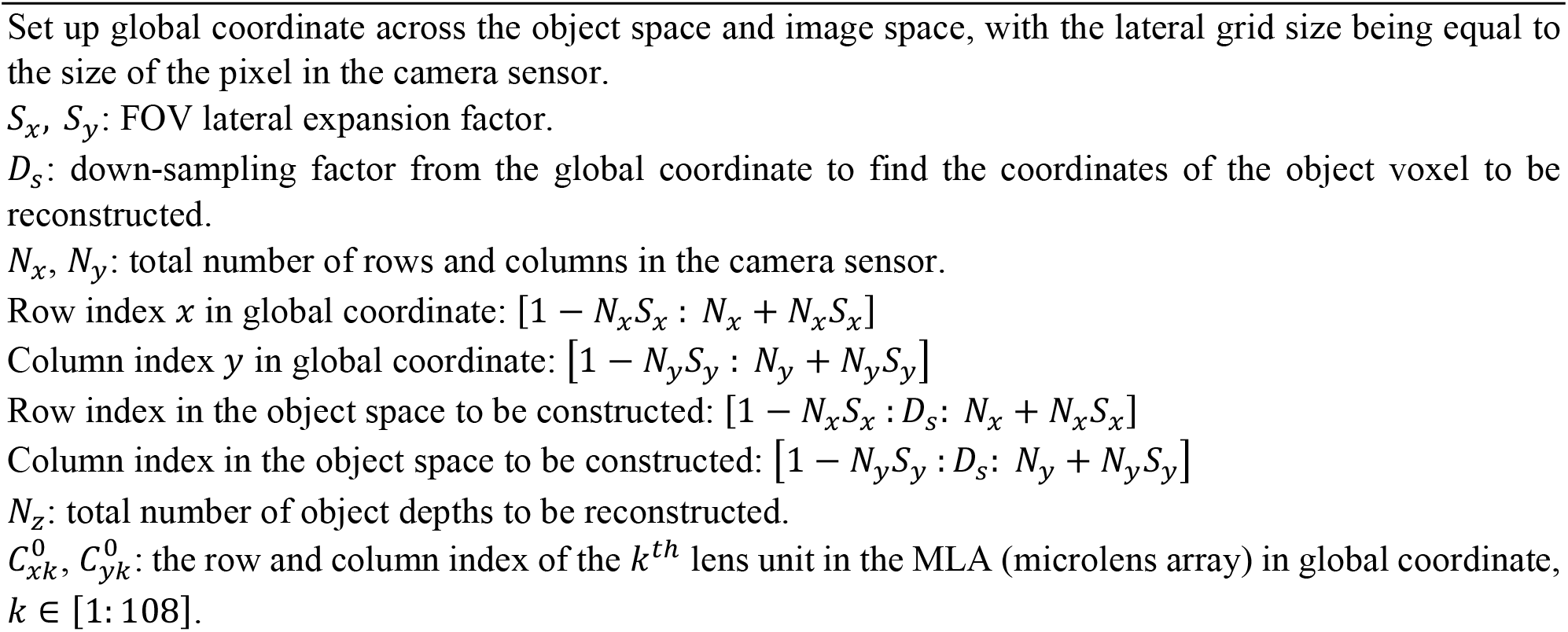
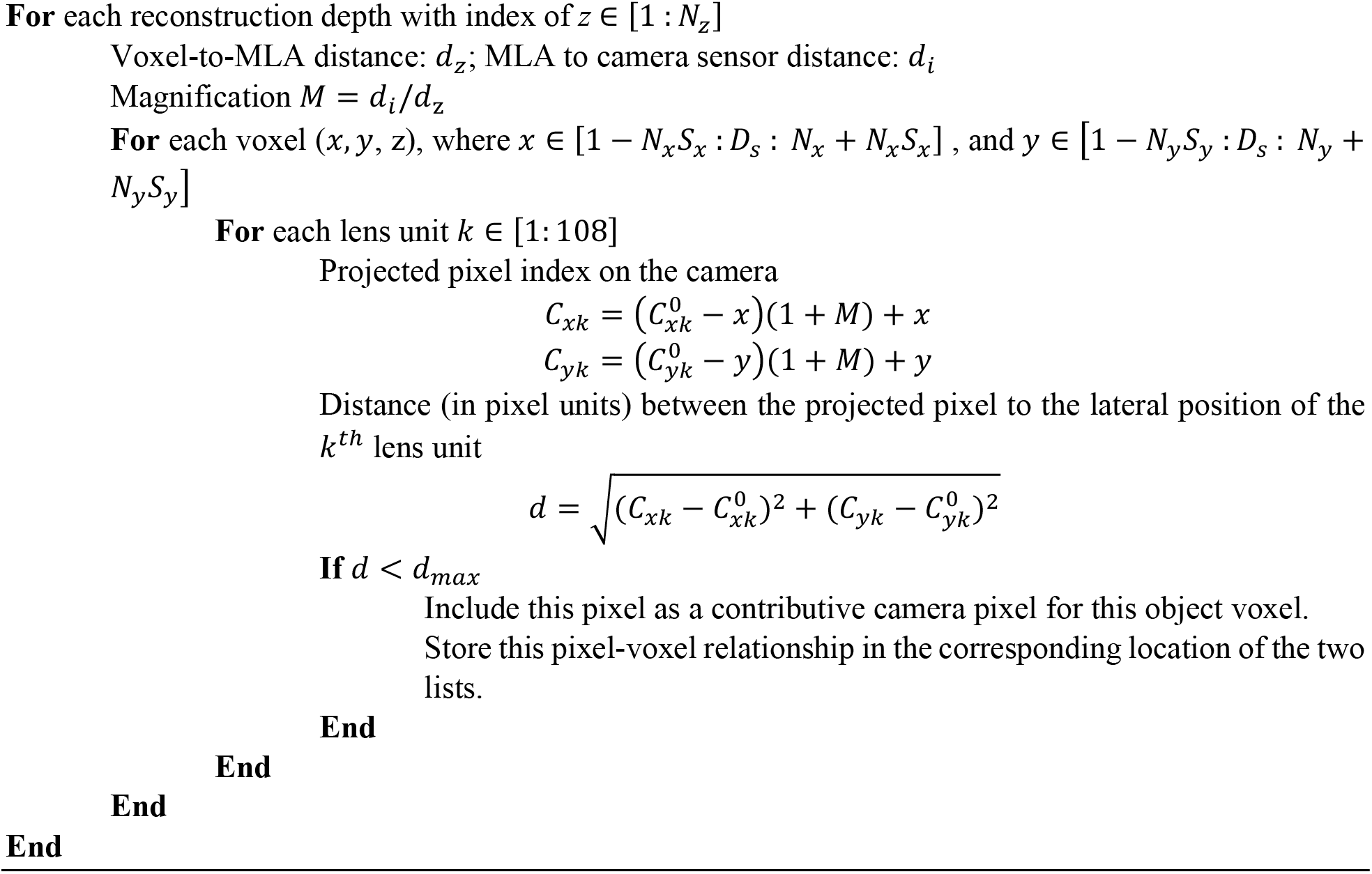

Using the two lists, we could conduct list-based RL algorithm. The algorithm follows the same procedure as the standard RL algorithm, except that the full system matrix to store the mapping relationship between the object voxel and camera pixel is replaced by the two sets of lists. We typically conducted 3∼5 iterations to achieve a good convergence.

The pseudo code below illustrates the list-based RL algorithm.

#### Algorithm 2: List-based RL

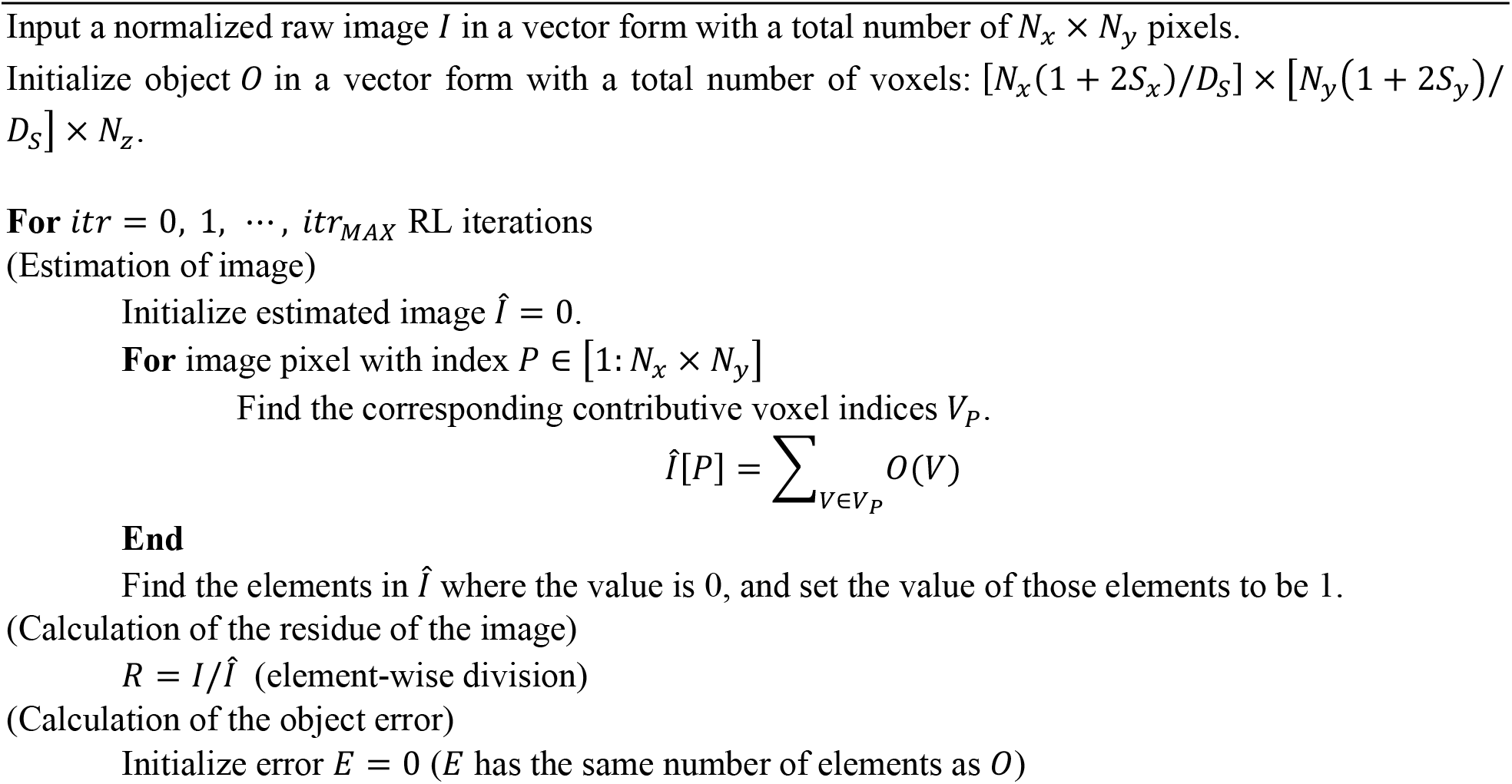
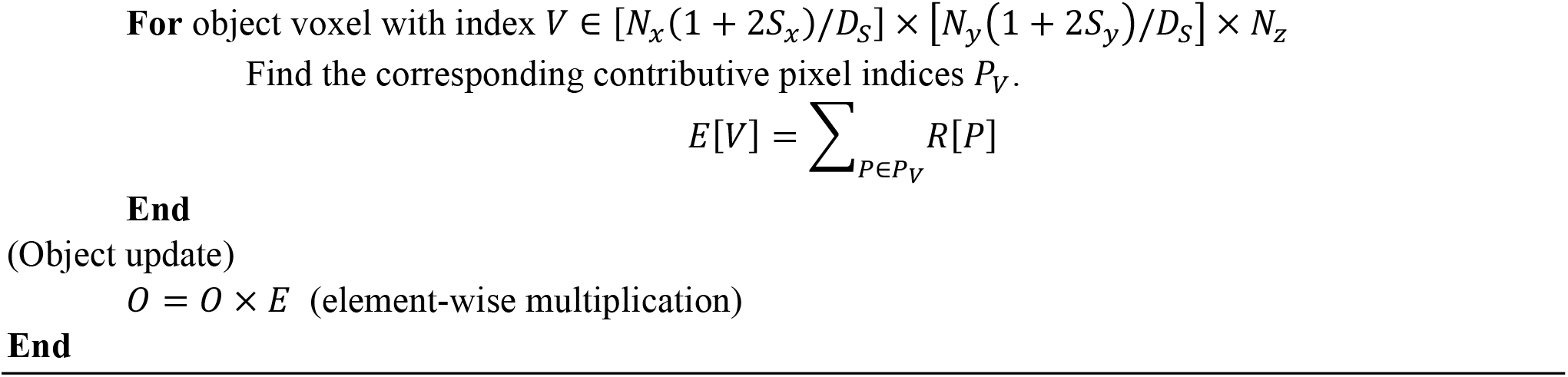

### Neural networks used to compare with multi-FOV ADMM-Net

We compared the performance of multi-FOV ADMM-Net against single-FOV Hadamard-Net and single-FOV ADMM-Net, as well as multi-Hadamard-Net and multi-Wiener-Net.

The single-FOV Hadamard-Net and single-FOV ADMM-Net have been described earlier. The Hadamard-Net is essentially the *X*_*o*_ initialization in stage 0 in ADMM-Net, without any subsequent stage for ADMM updates.

Both multi-Hadamard-Net and multi-Wiener-Net reconstruct multiple global FOVs and then fuse them together. The multi-Hadamard-Net shared a similar concept as Ref. ^28^. The image *b* is first transformed into 2D Fourier domain. An element-wise multiplication (i.e. Hadamard multiplication) between the 2D Fourier transform of *b* and multiple sets of 2D learnable kernels are then performed, respectively for both real components and imaginary components. Each resultant image, corresponding to each set of 2D learnable kernels, is then inverse Fourier transformed back to spatial domain. A U-Net then fuses these images together. Each of the kernel is initialized randomly. In the results shown in Fig. 1, we used 6 kernels.

The multi-Wiener-Net is adopted from Ref. ^26^. The original work requires PSF characterization across the entire FOV. To be consistent with other benchmarking methods, we initialized the PSF kernels randomly in multi-Wiener-Net. The noise level scalar was initialized as one. In the results shown in Fig. 1, we used 6 PSF kernels.

In all these methods, the raw measurement could be preprocessed by a denoise module, before processed by the neural networks.

### Evaluation metrics on reconstructions

We use both spatial domain and frequency domain metrics to evaluate the performance of the reconstruction algorithms. In spatial domain, we use peak-SNR (PSNR) to measure the peak signal quality, structure similarity index (SSIM) and correlation coefficient (CORRCOEF) to measure the spatial feature position similarity, and the learned perceptual image patch similarity (LPIPS) to measure perceptual level similarity, all versus the reference ground truth captured by a benchtop microscope.

In frequency domain, we followed Ref. ^32^ to analyze the frequency spectrum of the reconstructed images. After the 2D Fourier transform of the image into frequency domain, we created a 1D power spectrum by averaging the power amplitude at each frequency radius from the zero frequency center on the 2D image at the frequency domain. We then analyzed the high frequency components (upper 60% of the frequency spectrum) which correspond to fine features in the sample, in terms of mean energy (fMean) and standard deviation (fSTD). These metrics evaluate the similarity of features in the frequency domain between the reconstructed object and the reference ground truth. Importantly, they do not emphasize the spatial location of these features, and thus are resistant to shifts caused by alignment errors in the experiment.

### Preparation of scattering phantom sample

We prepared scattering phantom by mixing non-fluorescent microspheres in 1.18 µm diameter (Cospheric) in our customized thick phantom slide, composed of randomly distributed fluorescent microspheres/beads (12 µm diameter, Cospheric) in optical clear UV-light-curable polymer (Norland). The scattering mean free path was calculated by Mie scattering theory^33^:

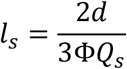

where *d* is the mean diameter of scatters, Φ is the volume fraction of scatters, and *Q*_*s*_ is the scattering efficiency factor calculated by Mie scattering calculator^34^. We calculated the mean free path following the Mie scattering formula. The non-fluorescent microspheres were weighted by analytical scales with precision of 0.1 mg, and powdered by ultrasonic machine before adding to the optical polymer to avoid clustering. The fluorescent beads were then added to the scattering medium and mixed uniformly with the polymer. The polymer was then poured to a 3D printed phantom slide and then cured by UV light.

### Mice surgery, experiment and image processing

All mouse experiments and housing procedures were conducted with the approval and under the guidance of University of California Davis Institutional Animal Care and Use Committee (IACUC). Wild-type C57BL/6J mice (2-5 months old) were injected with AAV1-hSyn-GCaMP6f into the right primary visual cortex and chronic craniotomy was performed to attach a circular 3.5 mm glass coverslip window for imaging. The center of the coverslip was aligned with the injection sites. A custom stainless steel headplate with a 7 mm hole in the center was attached to the skull surrounding the injection and craniotomy location through dental cement (MetaBond). The coverslip was secured with cyanoacrylate (3M VetBond) and further sealed with dental cement to cover the exposed bone in the center of the headplate, and to stabilize the headplate for head-fixed imaging. After a period of 3∼4 weeks, which allows for virus expression, awake imaging was performed. The headplate was fixed to custom posts on top of a 3D printed treadmill where the mouse could run freely. The neuronal activity was then imaged by the miniaturized lensless microscope positioned directly above the cranial window, with the excitation light delivered through the two fiber ports connected to the miniaturized lensless microscope.

The recorded raw video was preprocessed to find the pixels that had large variance in time, which presumably contained the neural activity of active neurons. This preprocessing stage was realized by an operation of standard derivation of difference-to-local-mean (STD-DLM): we first applied a box filter to each raw video frame to get the locally averaged brightness of each pixel, and then calculated the difference between the pixel value in the original raw frame and its local average; we then calculated a standard deviation for each pixel along the temporal dimension. This STD-DLM operation resulted in an image (termed as STD-DLM mask), which represented weights of local signal variation. We then used the STD-DLM mask to elementwise multiply with the raw videos. The temporal activities could then be enhanced and the background noise could be suppressed by the difference of the weights in different local areas. Each frame of the weighted raw video was then processed by the multi-FOV ADMM-Net for 3D reconstruction. We analyzed this time-series 3D reconstruction volume using two algorithms. First, we constructed a time-series standard-derivation project of the 3D volume, and used an iterative clustering algorithm^14^ to find spatially discrete clusters, which were putative neurons with strong activity. Second, we applied a constrained negative-matrix factorization algorithm (CNMF-E)^30^ to extract the spatial footprint and deconvoluted the activity of individual neurons for each reconstructed plane in the time-series 3D reconstructed volume. From the CNMF-E results, we could further select the neurons based on the clustering result. Such a selection enhanced the extraction fidelity of neurons.

## Supplementary Information

**Supplementary Figure S1.**
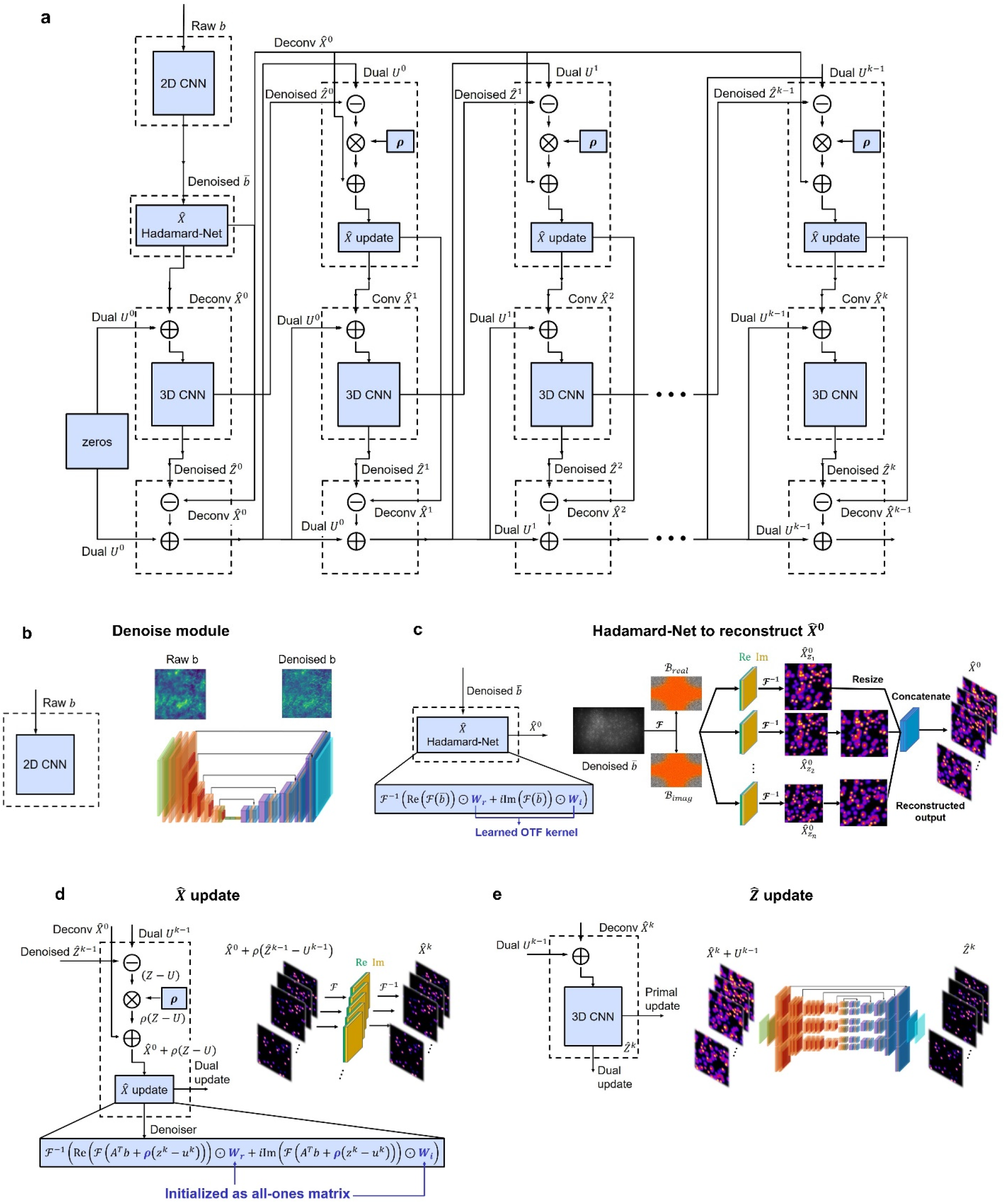
Architecture of ADMM-Net. (a) Expanded diagram of the ADMM-Net architecture in Figure 3. (b) Network architecture of the 2D U-Net used to preprocess the raw measured image to extract features and suppress the noise. (c) Network architecture of the Hadamard-Net (deconvolution module) to reconstruct 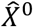. The preprocessed image from (b) is Fourier transformed to the frequency domain, with real and imaginary component separated. For each reconstruction depth, elementwise multiplication (i.e. Hadamard product) is performed between the learnable kernel weights and the real and imaginary part of the Fourier components of the raw image, respectively. The product is then inverse Fourier transformed to the spatial domain. Reconstructions at different depths are then resized to the same field of view size and concatenated into a 3D stack. The kernels are initialized from the pre-trained individual depth-specific 2D models. (d) Network architecture of 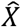 convolution module. Similar Hadamard products as (c) are applied at each depth and the outputs are concatenated into a 3D volume. The kernel weights are initialized as an all-one matrix. (e) Network architecture of the 3D U-Net denoiser, which update the regularized variable *Z*.

**Supplementary Figure S2.**
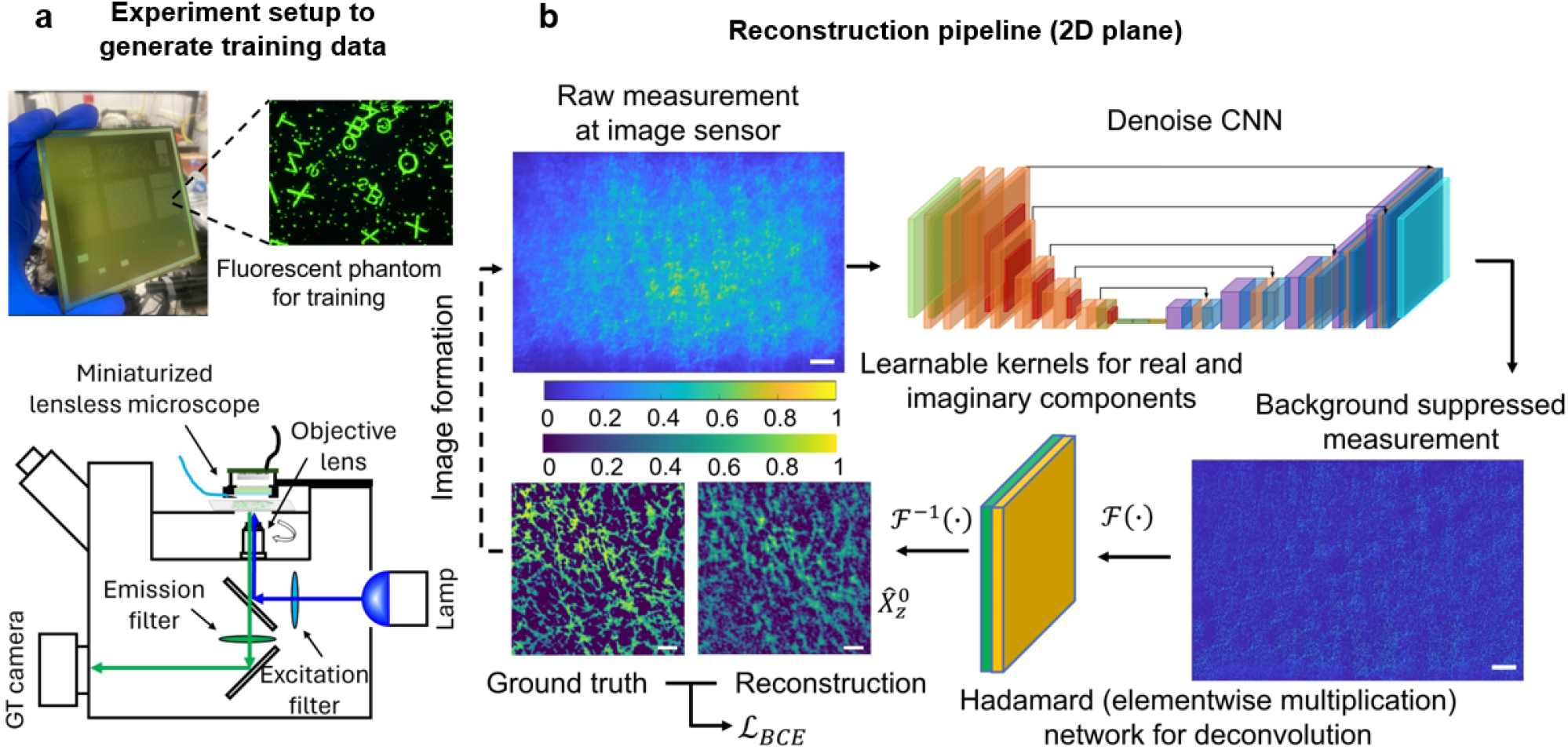
Training strategy using experimentally measured data. (a) Experimental setup used to generate training pairs in experiment. A customized photolithography mask with various features, painted with fluorescent spray (top), serves as samples. The sample is imaged by both a benchtop inverted microsope and the miniaturized lensless microscope DeepLeMiN, thus creating the training pairs. (b) Reconstruction pipeline for individual object depth. The pipeline contains two preprocessing modules including a noisy pixel removal module and a noise-suppression 2D U-Net module, and a Hadamard-Net to conduct a frequency domain deconvolution. The reconstructed results of each individual object depth are compared with the ground truth image from the benchtop inverted microscope. Scale bar, (b) 400 µm.

**Supplementary Figure S3.**
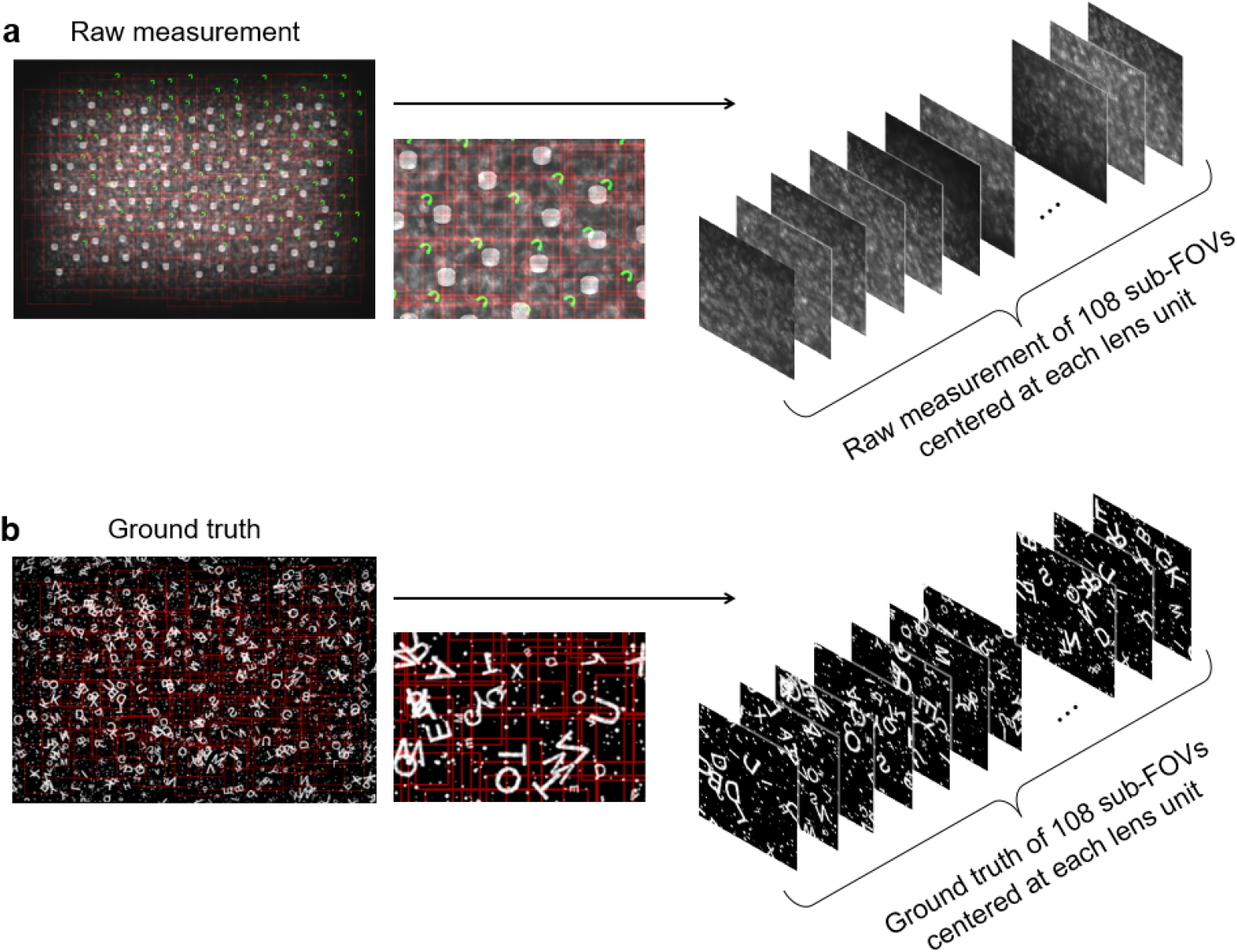
Local field-of-view (FOV) segmentation. (a) Raw image with sub-FOVs marked in red color. Each sub-FOV is centered at a lens unit on the microlens array. Each sub-FOV is rotated 180° to align with the ground truth patches. As there are 108 lens units, there are a total of 108 sub-FOVs. (b) Same as (a), but for ground truth images. Each sub-FOV from (a) and (b) forms a data pair for training.

**Supplementary Figure S4.**
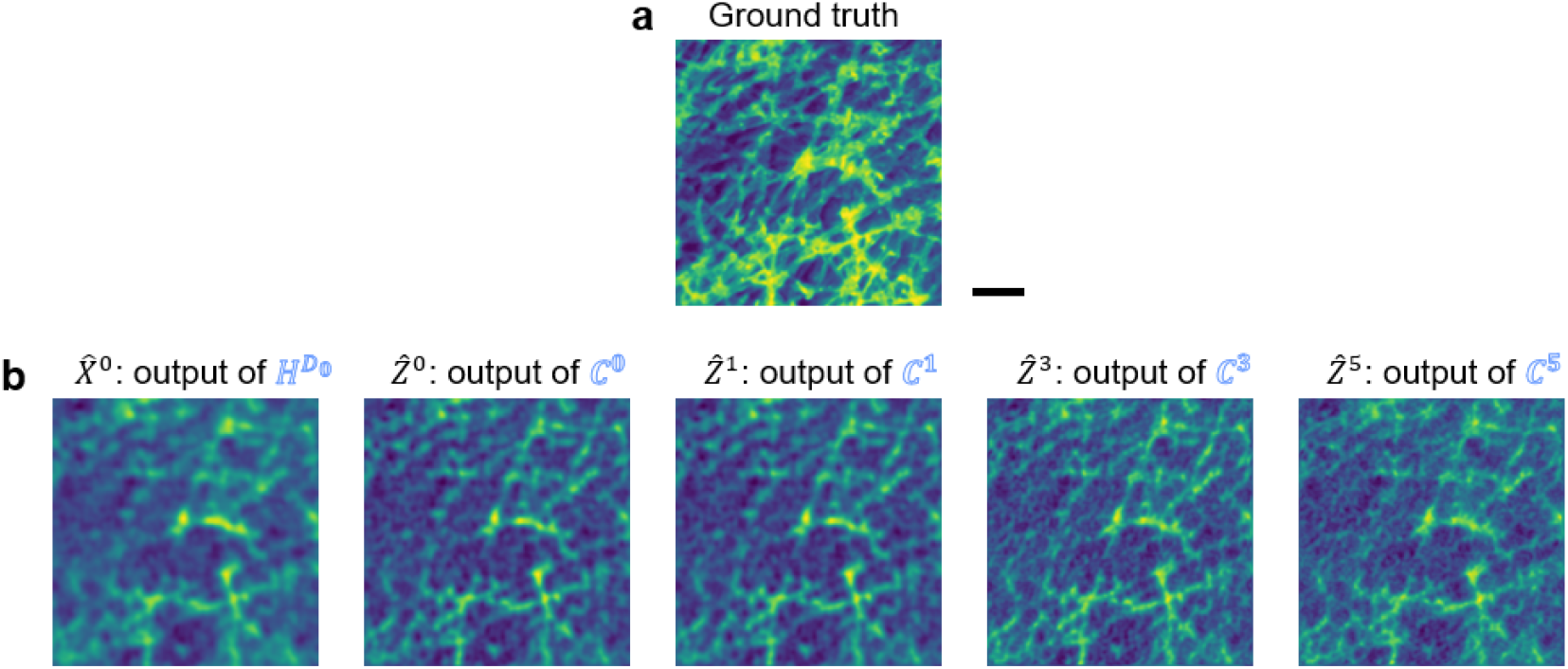
Comparison of outputs of stage 0 to stage 5 in ADMM-Net versus the ground truth. The comparison is for lens tissue sample stained with green fluorescent spray, in one out of the 108 fields of view. (a) Ground truth imaged by a benchtop microscope. (b) Reconstruction output from different stage of the ADMM-Net. As the stage number increases, the reconstruction has an improved quality, and gradually converges to the optimal results. Typically, the optimal reconstruction could be achieved within 3-5 stages in ADMM. Scale bar, 100 µm.

**Supplementary Figure S5.**
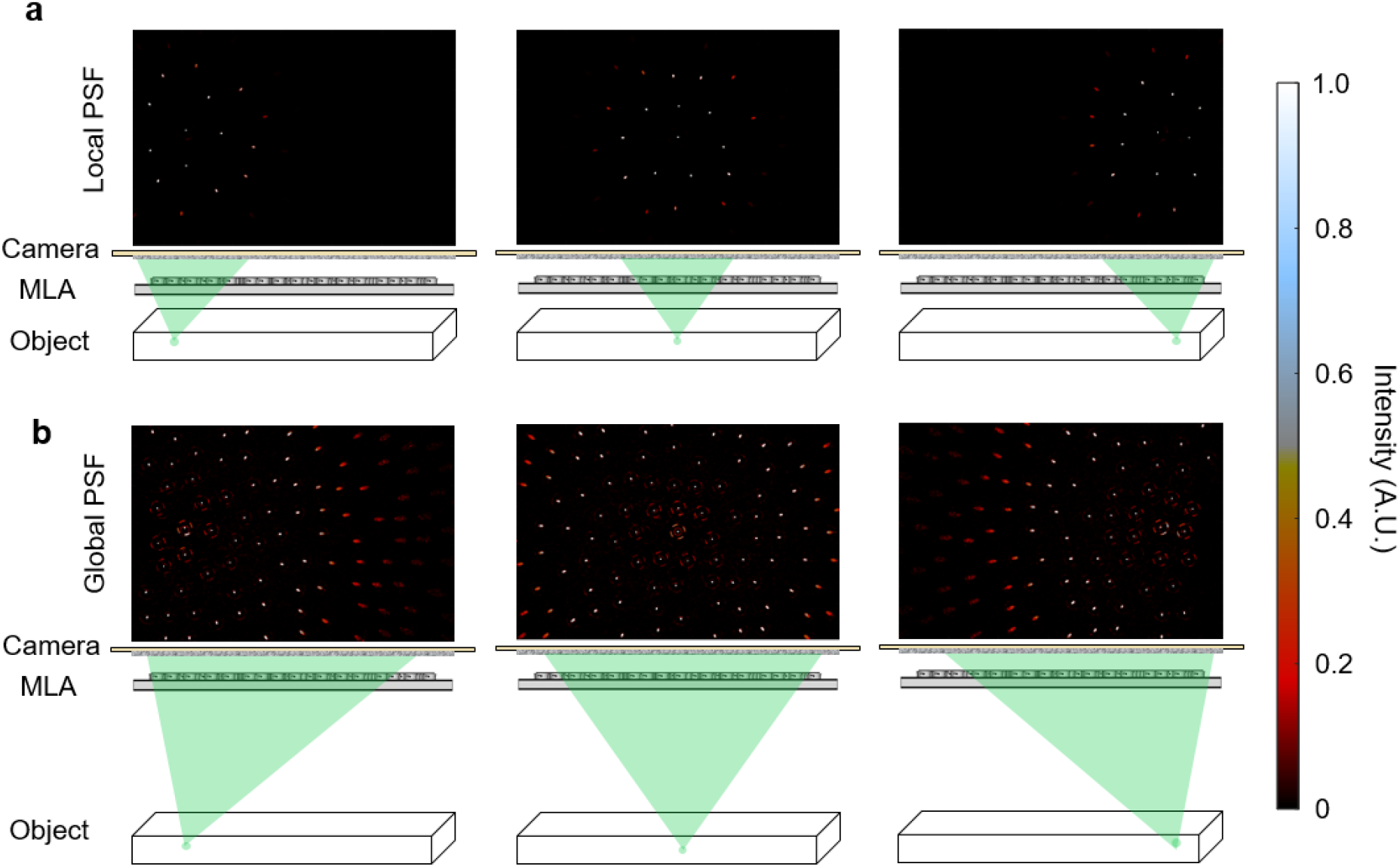
Comparison of local PSF/FOV and global PSF/FOV. We conducted ray tracing simulation (Zemax, Optics Studio) of the PSF of DeepLeMiN. (a) Concept of local PSF or local FOV. When the point source is close to microlens array, the PSF is local. The microlens units far away from the lateral position of the point source has negligible effect on the captured image. Each local region can be reconstructed independently and in parallel. Compared to reconstructing the entire FOV at once (i.e. global reconstruction), the local reconstruction strategy minimizes computational resources and achieves a faster reconstruction and a higher quality reconstruction. (b) Concept of global PSF or global FOV. When the point source is further away from the microlens array, the point source could be imaged by each microlens unit, and thus a global PSF pattern is formed on the image sensor. A global reconstruction strategy could be adopted. Note, the PSF could be spatial variant locally in (a), and globally in (b).

**Supplementary Figure S6.**
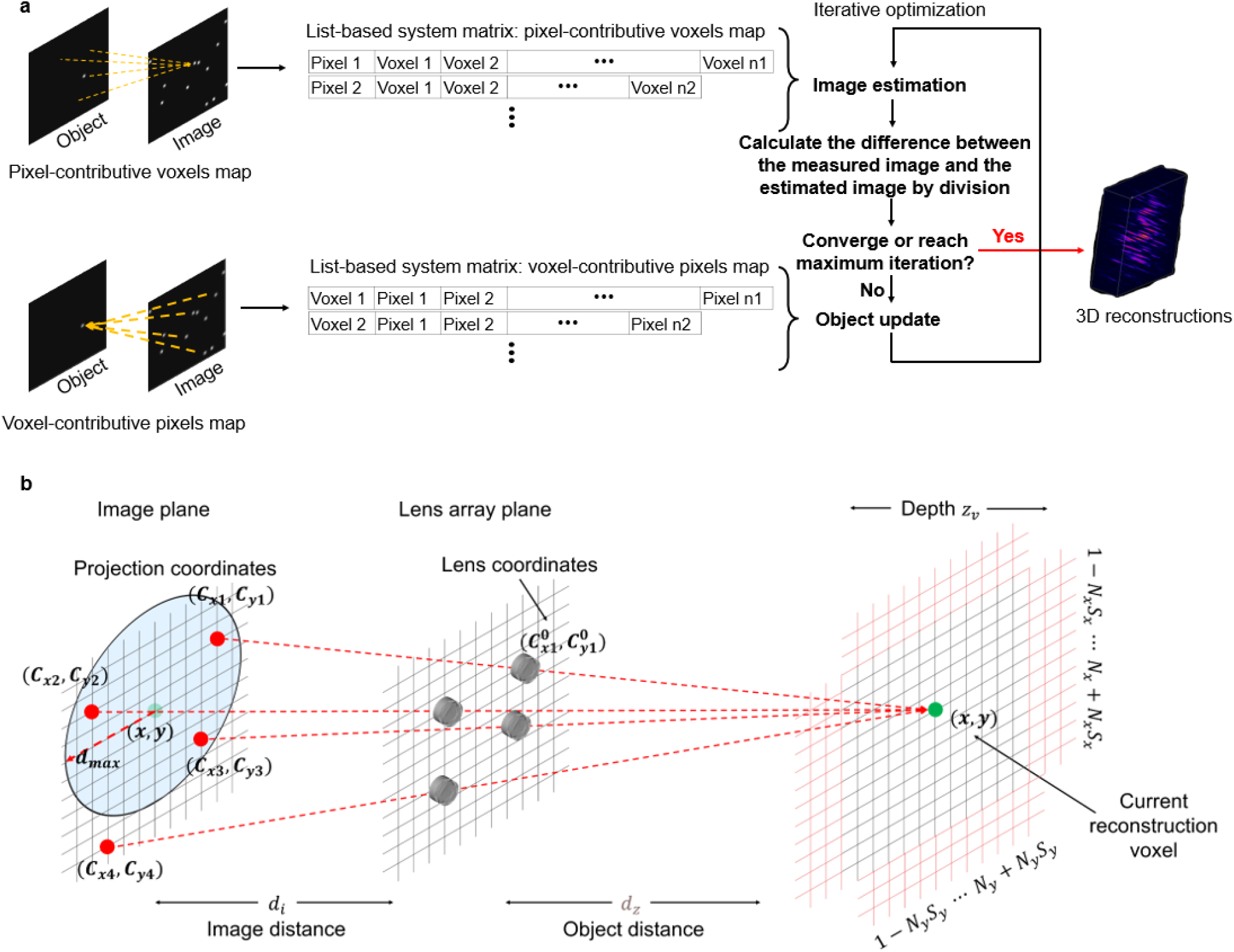
List-based Richardson-Lucy deconvolution algorithm. (a) List-based Richardson-Lucy iterative deconvolution algorithm for fast 3D reconstruction. The system matrix describing the mapping relationship between the object voxels and image pixels could be represented by two sets of lists. The image estimation and object update operate on the two sets of list, effectively reducing the complexity and required memory in the computation. (b) Illustration of how each object voxel is mapped to multiple camera pixels through the microlens units. A global coordinate is defined across the object space, microlens array and image plane. The row index *x* is [1 − *N*_*x*_*S*_*x*_ : *N*_*x*_ + *N*_*x*_*S*_*x*_], and the column index *y* is [1 − *N*_*y*_*S*_*y*_ : *N*_*y*_ + *N*_*y*_*S*_*y*_]. This illustration shows an object voxel (*x, y*) (axial index *z* is ignored here for clarity) is mapped to 4 pixels through a microlens unit. See Methods for more detailed explanation of the mapping.

**Supplementary Figure S7.**
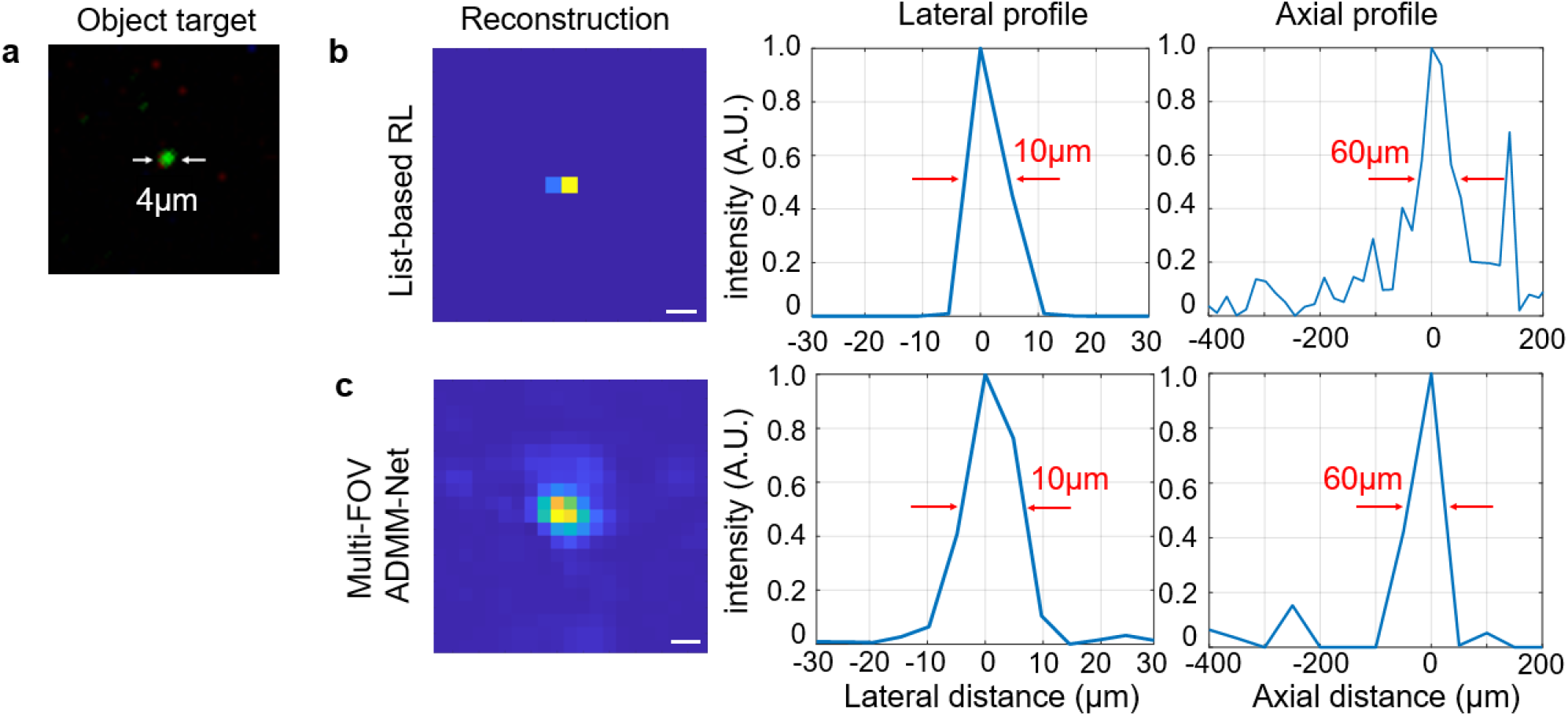
Comparison of the reconstruction results of a 4 µm point source using list-based RL and multi-FOV ADMM-Net reconstructions. (a) Object point source captured by a bench-top microscope. (b) Lateral 2D view (left), lateral line profile (middle) and axial line profile (right) of the reconstructed object, from the list-based RL algorithm. (c) Same as (b), but from a multi-FOV ADMM-Net. Scale bar, (b)-(c) 10 µm.

**Supplementary Figure S8.**
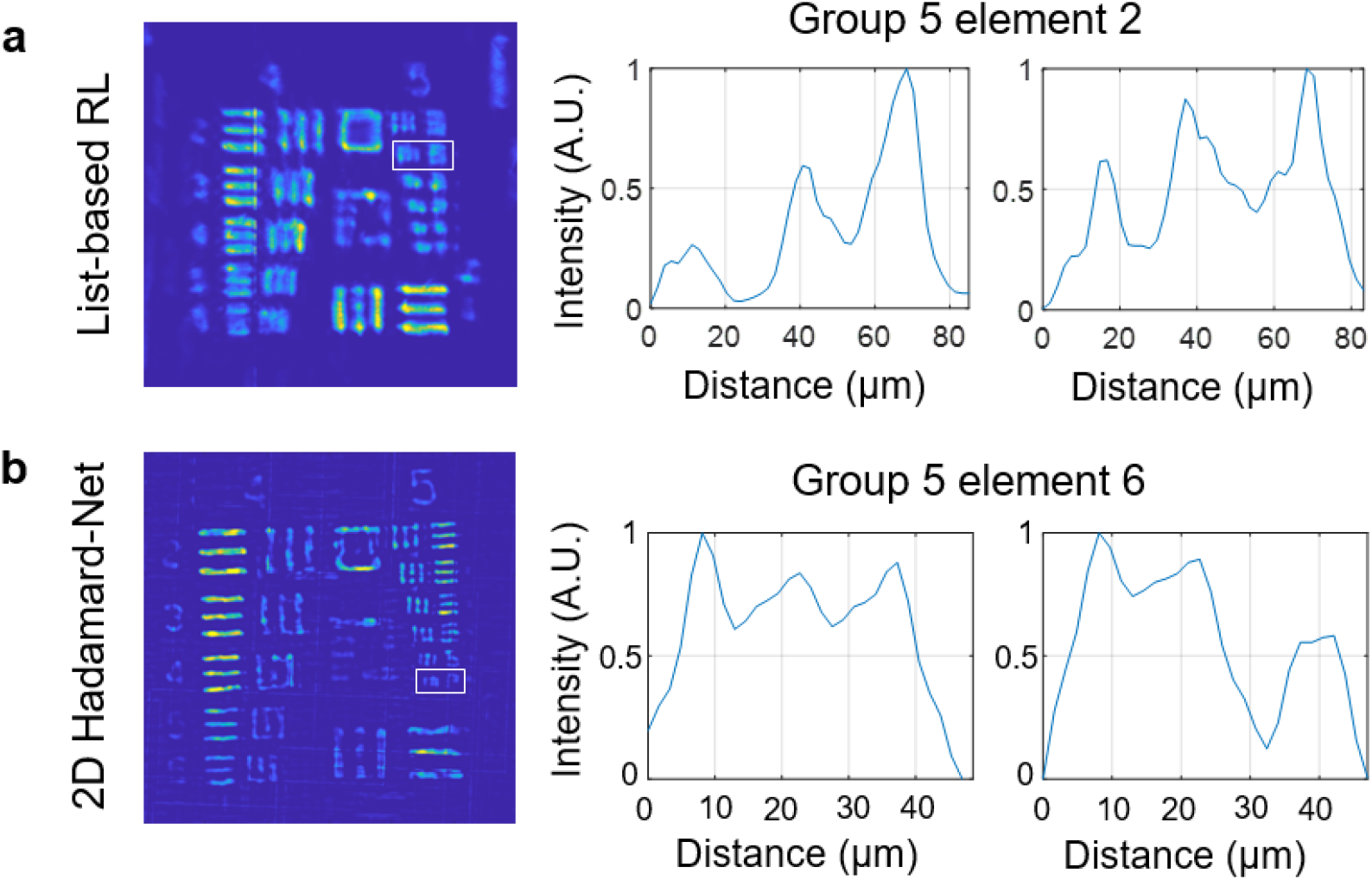
Comparison of the reconstruction results of USAF resolution target using list-based RL, and 2D Hadamard-Net reconstructions. (a) Lateral 2D view (left) and the line profiles of the group 5 element 2 (middle/right) of the reconstruction results from the list-based RL algorithm. (b) Lateral 2D view (left) and the line profiles of the group 5 element 6 (middle/right) of the reconstruction results from a 2D Hadamard-Net. For this sample, Hadamard-Net without ADMM update is sufficient to achieve high-quality reconstruction results.

**Supplementary Figure S9.**
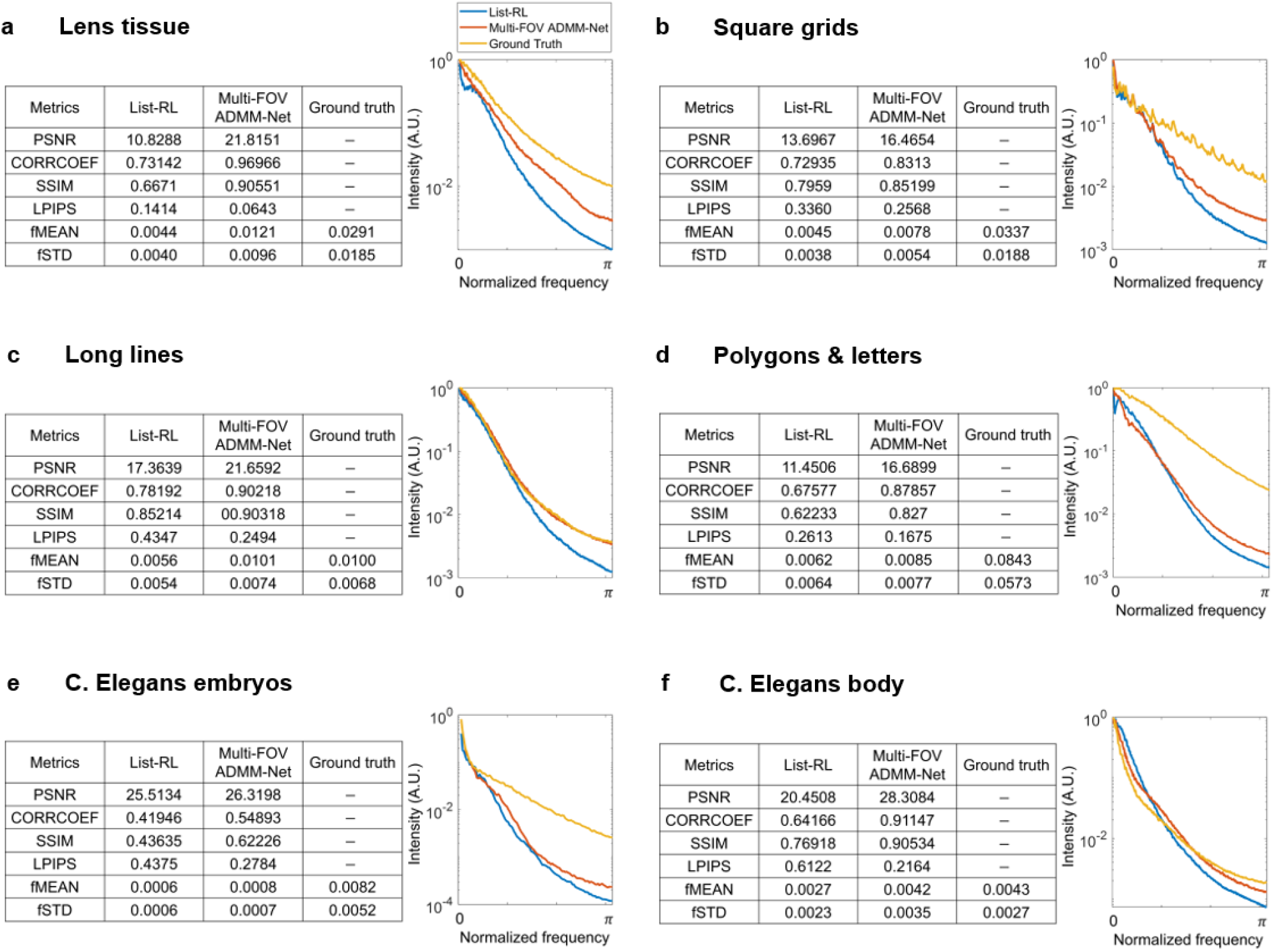
Spatial and frequency domain metrics to evaluate the reconstruction results of the various samples in Figure 1, from list-based RL algorithm and multi-FOV ADMM-Net. (a) Evaluation metrics of the reconstruction of lens tissue sample. Left, spatial metrics including peak-SNR (PSNR), correlation coefficient (CORRCOEF), structure similarity index (SSIM), learned perceptual image patch similarity (LPIPS) and frequency metrics including fMEAN and fSTD (Methods). Right, power spectrum. (b) Square grids sample. (c) Long lines samples. (d) Polygons and letters samples. (e) C. Elegans embryos. (f) C. Elegans body.

**Supplementary Figure S10.**
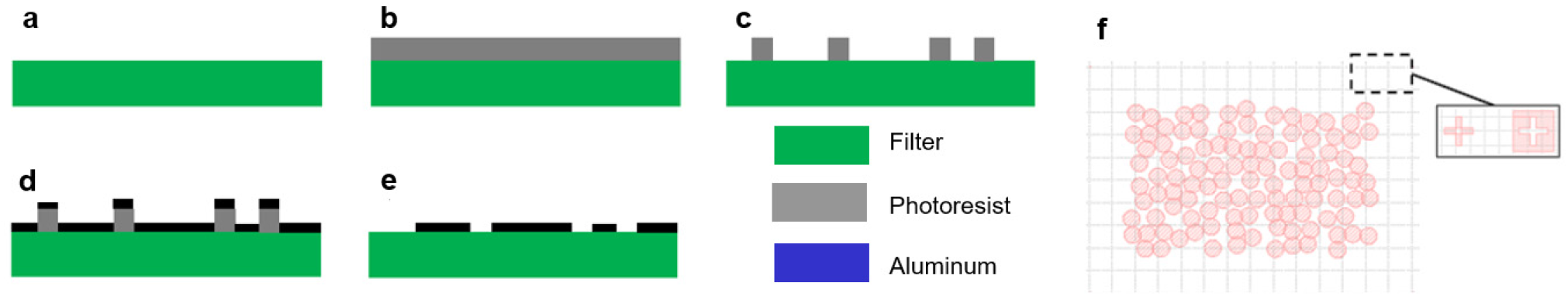
Fabrication flow to deposit aluminum on the filter. (a) Filter preparation. (b) Photoresist deposition. (c) Photolithography: exposure and development. (d) Metal deposition. (e) Liftoff. (f) Pattern of the mask for photolithography.

**Supplementary Figure S11.**
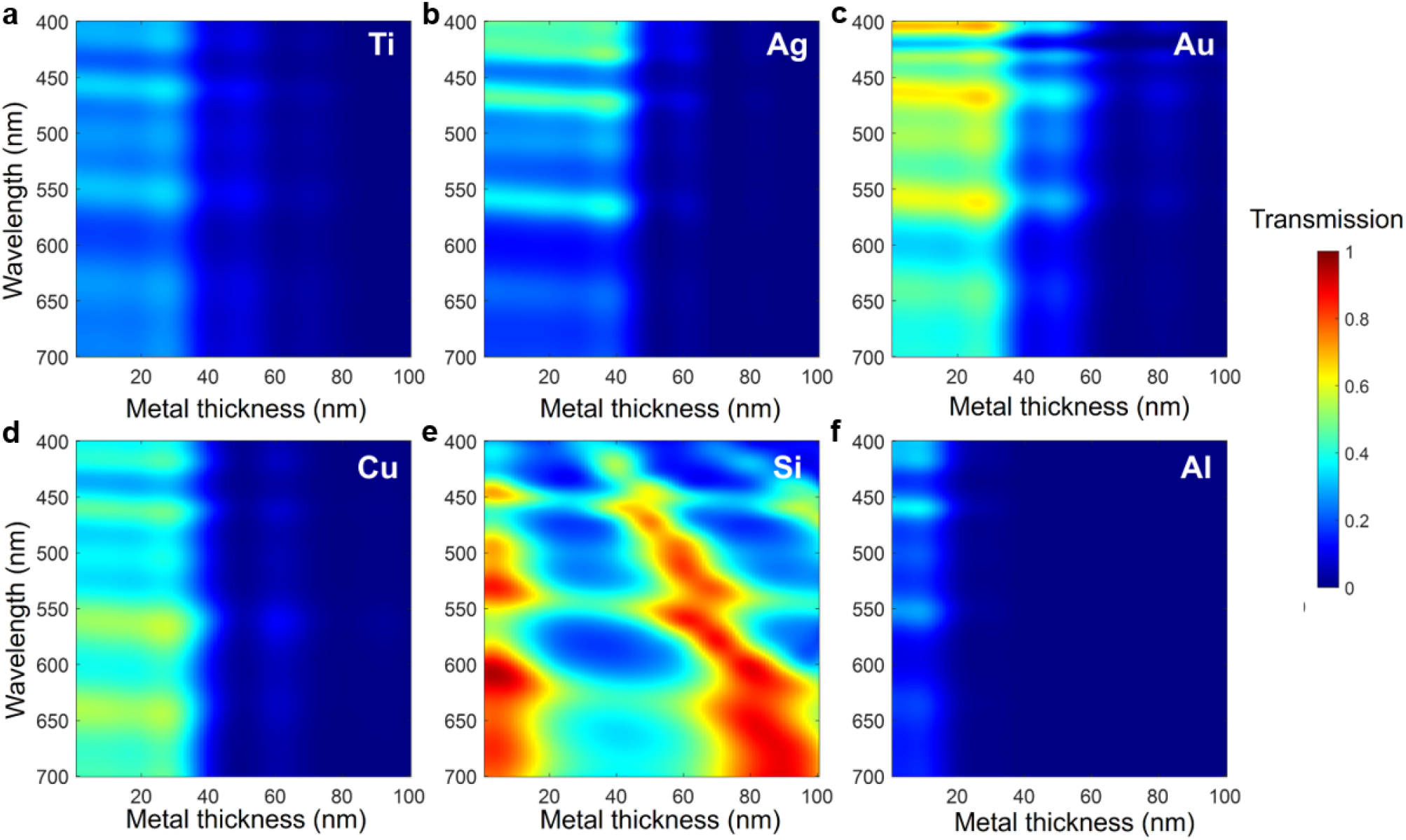
Transmission spectrum versus metal thickness, simulated for different metal through finite-difference time-domain (FDTD) method. The simulation results show that aluminum has best reflectivity to block the background light that goes into gaps between the microlens units on the array. (a)-(f) Titanium (Ti), silver (Ag), gold (Au), copper (Cu), silicon (Si) and aluminum (Al).

